# Diffusion Tensor Model links to Neurite Orientation Dispersion and Density Imaging at high b-value in Cerebral Cortical Gray Matter

**DOI:** 10.1101/441659

**Authors:** Hikaru Fukutomi, Matthew F. Glasser, Katsutoshi Murata, Thai Akasaka, Koji Fujimoto, Takayuki Yamamoto, Joonas A. Autio, Tomohisa Okada, Kaori Togashi, Hui Zhang, David C. Van Essen, Takuya Hayashi

## Abstract

Diffusion tensor imaging (DTI) and neurite orientation dispersion and density imaging (NODDI) are widely used models to infer microstructural features in the brain from diffusion-weighted MRI. Several studies have recently applied both models to increase sensitivity to biological changes, however, it remains uncertain how these measures are associated. Here we show that cortical distributions of DTI and NODDI are associated depending on the choice of b-value, a factor reflecting strength of diffusion weighting gradient. We analyzed a combination of high, intermediate and low b-value data of multi-shell diffusion-weighted MRI (dMRI) in healthy 456 subjects of the Human Connectome Project using NODDI, DTI and a mathematical conversion from DTI to NODDI. Cortical distributions of DTI and DTI-derived NODDI metrics were remarkably associated with those in NODDI, particularly when applied highly diffusion-weighted data (b-value =3000 sec/mm^2^). This was supported by simulation analysis, which revealed that DTI-derived parameters with lower b-value datasets suffered from errors due to heterogeneity of cerebrospinal fluid fraction and partial volume. These findings suggest that high b-value DTI redundantly parallels with NODDI-based cortical neurite measures, but the conventional low b-value DTI does not reasonably characterize cortical microarchitecture.

## 1. Introduction

The diffusion motion of water molecules in brain tissue is affected by the local microarchitecture, including axons, dendrites and cell bodies. Diffusion tensor imaging (DTI) is a well-established model that describes Gaussian properties of diffusion motion in a fibrous structure like brain white matter from diffusion-weighted MRI (dMRI)^1,2^, and is widely used for inferring the microstructural changes related to plasticity and diseases (for review, Johansen-Berg and Behrens, 2013)^3^. In most cases, summary parameters of DTI, fractional anisotropy (FA) and mean diffusivity (MD), have been studied based on dMRI data acquired with low b-value (b-value less than 1000), however, these parameters have not been shown to be specific to underlying microstructural features of axons and dendrites (collectively referred to as neurites) and are often sensitive to tissue compartments other than neurites^4^.

One recent advance for estimating the microstructural complexity of brain tissue using dMRI is the Neurite Orientation Dispersion and Density Imaging (NODDI)^5^. NODDI models dMRI signals by combining three tissue compartments: neurites, extra-neurites, and cerebro-spinal fluid (CSF), each with different properties of diffusion motion, and enables in vivo estimation of a neurite density index (NDI) and an orientation dispersion index (ODI), as well as a volume fraction of isotropic diffusion (V_iso_). NODDI requires dMRI data to be scanned with multiple b-values (e.g. b=700 and 2000 sec/mm^2^) and relatively higher number of diffusion gradient directions (e.g. >90 directions over two b-shell) as compared with DTI^5^. The NDI estimates the volume fraction of neurites, including both axons and dendrites, whereas the ODI estimates the variability of neurite orientation: ranging from 0 (all parallel) to 1 (isotropically randomly oriented). NODDI has already been applied to many studies because of their feasibility. The variation of NODDI estimates in white matter have been related to aging^6–11^ and neurologic disorders^12–14^. Of note, NODDI has proven to be useful for gray matter neurite changes as reported in several clinical studies, e.g. in patients with IFN-α-induced fatigue^15^, Wilson’s disease^16^, cortical dysplasia^17^, aging^18^, and schizophrenia^19^. We recently optimized NODDI for cortical gray matter^20^, finding that the NODDI reflects neurobiology of cortical microarchitecture – cortical distribution of NDI is closely related to cortical myelin^21^ and ODI is associated with cortical organization of radial/horizontal fibers^22,23^. Although there is recent debate about oversimplified assumptions in NODDI such as uniform diffusivity^24^, it is of note that accumulated histological evidence indicates that NDI and ODI of neural tissues are reasonably representing histology-based neurite density^25^ and orientation dispersion^25–28^, respectively.

Recently, there is accumulating evidence of combined DTI and NODDI analysis in clinical studies. Those performed both DTI and NODDI in the pathological cortex all showed opposite changes between MD and NDI in Parkinson’s disease^29^, multiple sclerosis^25^, and stroke^30^. Our previous study also revealed that strong relationships between NODDI and DTI parameters in the cortex of healthy subjects, in particular, NDI and 1/MD were very highly correlated (R=0.97)^20^ when used three-shell dMRI data including high b-value, but not so highly correlated when used low b-value data. On the other hand, in vitro study showed that slower-decaying component found by high b-value dMRI signals were originated from intra-neurite water^31^, thus suggesting that MD obtained at high b-value is specifically reflecting neurite properties. High b-value DTI in clinical studies also implicate higher sensitivity to neurobiological changes than low b-value, e.g. the contrast between gray/white matter^32^, ischemic/hypoxic changes in the gray matter^33^, white matter disintegrity in schizophrenia^34^ and maturation in juveniles^35^. However, there is no consensus how DTI is associated with NODDI parameters, and how it is dependent on the b-shell scheme.

In this study, we investigated how DTI and NODDI parameters are related with each other in cortical gray matter in healthy subjects. The measures were correlated by two methods and also analyzed by utilizing a recently derived mathematical function, which converts DTI to NODDI parameters^24,36^. We used the preprocessed dMRI data from Human Connectome Project (HCP), and estimated b-shell scheme dependency of the relationship between DTI and NODDI parameters. Since the function relies on the assumption that CSF compartment (V_iso_) is negligible in the tissue^24,36^, we also estimated V_iso_ in the cortex and the white matter based on previous work^20^ and estimated effect on the partial voluming and the relationship between DTI and NODDI parameters. We performed simulation analysis in terms of b-value, proportion of CSF signal. Our main purpose is to highlight the neurite properties in the specific subtype of brain, cortical gray matter, in healthy subjects, and investigate how DTI specifically represent the cortical NODDI metrics. We also review the past literature which applied NODDI and DTI in vitro and in vivo dMRI studies and discuss the potential interpretability of DTI.

## 2. Materials and Methods

In order to comprehensively investigate the relationship between NODDI and DTI in cortical gray matter, we examined whether NODDI parameters can be accurately estimated from DTI using their mathematical relation. Publicly available MRI data from 456 healthy subjects (aged 22-35 years) the HCP (https://www.humanconnectome.org/) were used. In particular, dMRI datasets with different b-shell structures were analyzed to investigate how the b-shell scheme affects the relationship between two diffusion model. To clarify why their relationship depends on the diffusion scheme, we also performed simulation analysis addressing several error sources such as CSF signals in dMRI data and partial volume effects. Data analyses were performed at RIKEN, and the use of HCP data in this study was approved by the institutional ethical committee (KOBE-IRB-16-24).

### 2.1.1 Subjects and dMRI datasets

We used the ‘S500 Release Subjects’ dataset from the publicly available HCP dataset, including high-resolution structural images (0.7-mm isotropic T1w and T2w images,^37^ and dMRI data (1.25-mm isotropic resolution)^38^. The dMRI data obtained with TR=5520ms and TE=89.5ms included 270 volumes with 90 volumes for each of the three shells of b-values (b=1000, 2000 and 3000 s/mm^2^) in addition to 18 non-diffusion weighted (b=0 s/mm^2^) volumes. From this dataset, 456 healthy subjects (age, 22-35 years) scanned with a complete dataset of 270 volumes were chosen, and 49 subjects were excluded based on incomplete dMRI scans. In our previous study, NDI and the reciprocal of MD (1/MD) showed very similar surface distributions when all of the dMRI data were used, but they did not show similar distributions when only a single shell of b=1000 dMRI data was used^20^. Therefore, we hypothesized that the relationship between DTI and NODDI may differ depending on the b-shell scheme of dMRI data. To address this, datasets with different b-shell schemes were used for analysis (Table 1), i.e. for each subject, seven types of b-shell datasets were derived from dMRI data as follows: three one-shell datasets using b=0 volume and any one of b=1000, 2000, or 3000 volume; three two-shell datasets using b=0 images and any two of b=1000, 2000, or 3000 volume; and a three-shell dataset using all images.

**Table 1.**
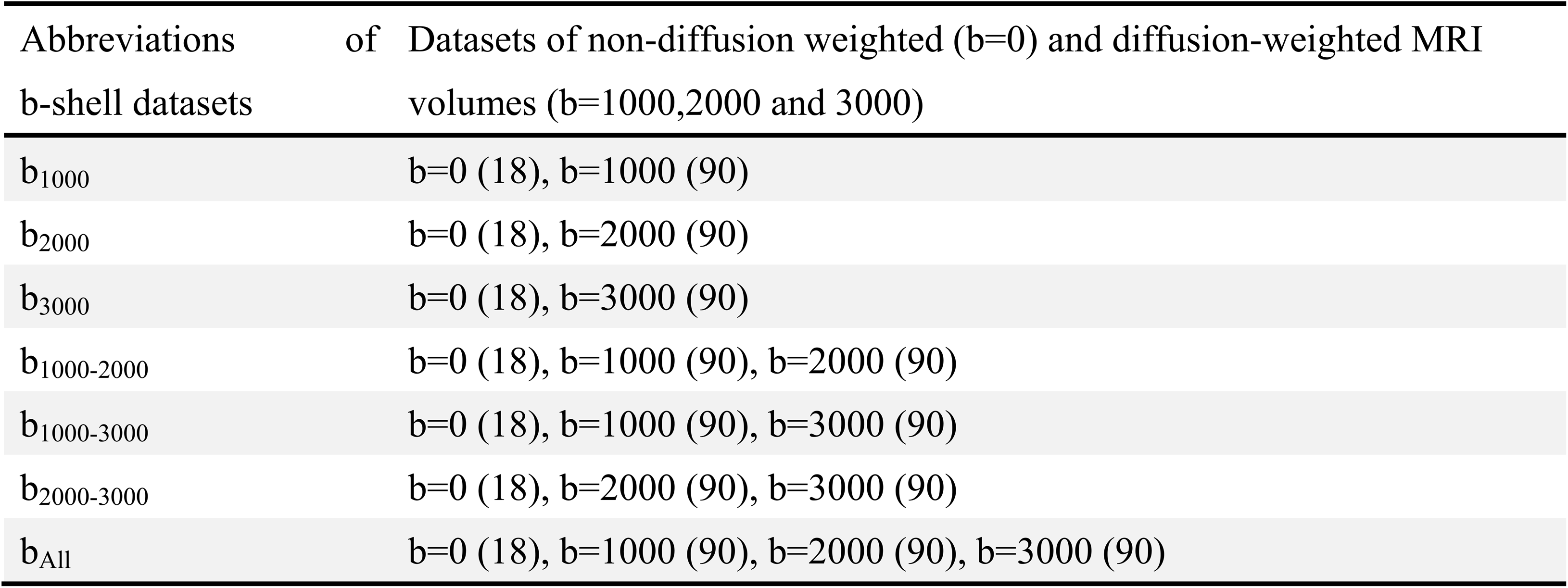
The table lists abbreviations of b-shell datasets used in the main text and corresponding datasets of dMRI in different b-shell schemes. The numbers in parentheses indicate the number of b0 volumes with repeatedly obtained for b=0 volume or diffusion weighted directions with different b-vectors (or directions of diffusion-weighted gradient) for each of the b=1000, 2000 and 3000 shells.

### 2.1.2 Calculation of the cortical surface map of original NODDI and DTI-derived NODDI parameters

The DTI estimates (FA and MD) were calculated using each dataset of dMRI and the dtifit diffusion tensor modeling tool in Functional Magnetic Resonance Imaging of the Brain Software Library (FSL) 5.09 (http://www.fmrib.ox.ac.uk/fsl). This linear DTI model was applied not only to a single-shell ‘standard’ b=1000 data but also to high b-value and multi-shell dMRI datasets. The primary reason of ‘forced’ fitting of DTI to such data was that we unexpectedly found correspondence of DTI metrics with those of NODDI in our previous paper. While it is known that these high b-value/multi-shell data take non-Gaussian distribution thus are more appropriate to apply a non-linear model like diffusion kurtosis imaging (DKI)^39^, there have been also a few reports that high b-value DTI sensitively detect the brain tissue pathologies^32–35^. The diffusion data were also fitted to the NODDI model using the optimized value of d_//_ and Accelerated Microstructure Imaging via Convex Optimization (AMICO) 1.0^40^, which re-formulates the original NODDI model as a linear system and shortens the calculation time. The value of d_//_ was optimized for the cerebral cortex (1.1 × 10^-3^ mm^2^/s) and changed from the default value (1.7 × 10^-3^ mm^2^/s)^20^, because we are interested in the cerebral cortical gray matter. We used default values of regularization (λ=0.001 and γ=0.5) for AMICO.

The parameters of the original NODDI model (NDI_ORIG_ and κ) and the DTI model (FA and MD) were mapped onto the cortical surface, as described previously^20^. Briefly, the algorithm for surface mapping identifies cortical ribbon voxels within a cylinder orthogonal to the local surface for each mid-thickness surface vertex on the native mesh and weights them using a Gaussian function (FWHM= ∼4 mm, σ=5/3 mm), which reduces the contribution of voxels that contain substantial partial volumes of CSF or white matter^21^. The ODI in the original NODDI (ODI_ORIG_) was calculated using the surface metric of κ and equation (5). The maps of DTI-derived NODDI parameters (NDI_DTI_ and ODI_DTI_) were calculated by converting from DTI maps to NODDI maps based on the mathematical relation (Fig. 1 and Supplementary text 1.2). The surface maps were resampled using MSMAll surface registration^41–43^ and onto the 32k group average surface mesh. For surface-based analysis, we used Connectome Workbench (https://github.com/Washington-University/workbench, Marcus et al., 2013). The scripts used in this manuscript are available from NoddiSurfaceMapping (https://github.com/RIKEN-BCIL/NoddiSurfaceMapping).

**Figure 1.**
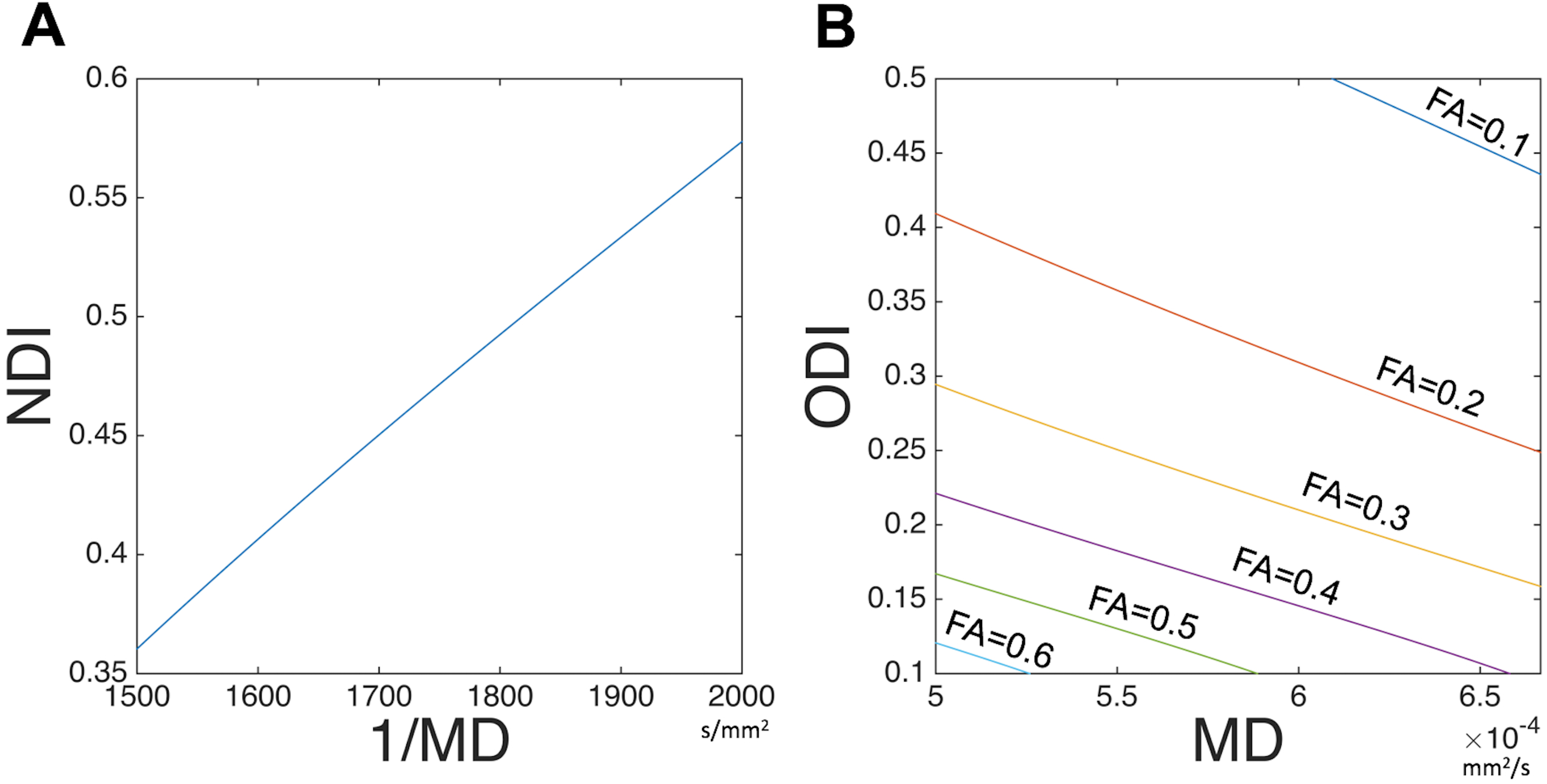
Relationships of DTI and NODDI when assumed non-CSF compartment. The equations for DTI-derived NODDI calculation (Eq. 2-5) and d// =1.1×10-3 mm2/s (optimized for gray matter) were used to simulate relationships between (A) Neurite density index (NDI) vs inversed mean diffusivity (1/MD), over the range of MD= 1500 to 2000 s/mm2, and (B) orientation dispersion index (ODI) vs MD when fractional anisotropy (FA) ranged from 0.1 to 0.6. Details of derivation of mathematical function of DTI and NODDI are described in Supplementary text 1.2.

### 2.1.3 Statistical analysis

Surface maps of NDI_ORIG_, ODI_ORIG_, V_iso_, NDI_DTI_ and ODI_DTI_ using each dataset were averaged across subjects and parcellated using the HCP’s multi-modal cortical parcellation (HCP_MMP1.0 210P MPM version)^41^. The mean value of each measure for each of the 180 parcels per hemisphere was calculated. NDI_ORIG_ and ODI_ORIG_ calculated using all the dMRI data were considered ‘a gold standard’ reference. To investigate the linear relationship between DTI-derived NODDI parameters and the original NODDI parameters, the correlations between each parcellated surface map (NDI_ORIG_, ODI_ORIG_, NDI_DTI_ and ODI_DTI_) and the reference in each subject were calculated using Pearson correlation analysis. To investigate whether DTI-derived NODDI parameters are biased, Bland-Altman analysis was performed in each dataset^45^. Briefly, Bland-Altman analysis is a method to confirm the presence or absence and degree of systematic bias visually by creating a scatter diagram (Bland-Altman plot), which is created by plotting the difference between two pairs of measured values on the y axis and the average value of the two measured values on the x axis.

Since the quality of the NODDI estimates depends on the image quality and preprocessing, we estimated the practical quality by the temporal signal-to-noise ratio (tSNR) of preprocessed b=0 volumes and removed 29 surface parcels with tSNR<17 from the analysis. Therefore, a total of 331 parcels in the whole cortex were used for the analysis. The cutoff was determined empirically in our previous study^20^.

### 2.2 Simulation for the effect of heterogeneity in CSF volume fraction on parameters of NODDI, DTI and DTI-derived NODDI

Since correlations and biases between original NODDI and DTI-derived parameters were dependent on the presence of high b-value data (b=3000 s/mm^2^) in the datasets (see section 3.1), simulations were performed to clarify whether and how potential sources of error can explain our findings of cortical DTI-derived parameters. One of potential sources is the amount of CSF compartment (V_iso_) in the voxel, which may be sum of intra-tissue CSF volume and partial voluming of CSF in the subarachnoid space (see also Supplementary text 2, Fig. S1). This compartment is not considered in the DTI or assumed to be zero in the DTI-derived NODDI calculation. The various levels or ‘heterogeneity’ of CSF volume fraction in the cortical voxel can cause errors at various level and could result in biases of the cortical distribution. Although actual heterogeneity of CSF volume fraction in the cortex cannot be measured in vivo, the simulation for the error sensitivity of diffusion measures to varying level of CSF volume fraction may give some insights. Another source of the error may be an interaction between heterogeneity CSF volume fraction and b-shell scheme of the data, since low b-value dMRI data may contain more CSF signal than high b-value dMRI data. Therefore, we tested how variable level of CSF and b-shell scheme can cause changes in the parameters of the original NODDI, DTI and DTI-derived NODDI in comparison with those calculated in reference condition of NDI=0.25, ODI=0.30 and V_iso_=0.1, the mean value of cortex^20^. The values of V_iso_ were varied from 0 to 0.6 (i.e. ΔV_iso_ from −0.1 to 0.5 in reference to V_iso_=0.1) with an interval of 0.1. Parameters of NODDI was calculated using the simulated three-shell dataset (b_All_) and those of DTI and DTI-derived NODDI was calculated using any of seven b-shell datasets (Table 1). The simulation data was created based on the mathematical equations and derivation described in the Supplementary text 3.1. To confirm the specificity of this findings, we also performed another simulation in which ‘homogeneous’ but small CSF volume fraction was assumed (Supplementary Text 3.2). The bias of DTI-derived NODDI and DTI parameters were also assessed by Bland-Altman analysis. (Supplementary text 3.2).

## 3. Results

### 3.1. Cortical maps of DTI-derived NODDI parameters using in vivo dMRI data

When the three-shell dataset (b_All_) in 456 subjects of HCP data were used in the original NODDI, the cortical map of neurite density (NDI_ORIG_) showed high intensity in the primary sensorimotor, visual, auditory cortices as well as the middle temporal (MT) area (Fig. 2A), while ODI_ORIG_ showed high intensity in the primary sensory, visual and auditory areas (Fig. 3A), as we reported previously^20^. Moreover, consistent with our previous study^20^, the cortical distribution of the NDI_ORIG_ was quite similar to that of the myelin map based on the T1w and T2w images, while the distribution of ODI_ORIG_ showed high contrast in the ‘granular cortex’ of von Economo and Koskinas ^23^, where cortical thickness is low and both radial and horizontal fibers are intermingled^20^.

**Figure 2.**
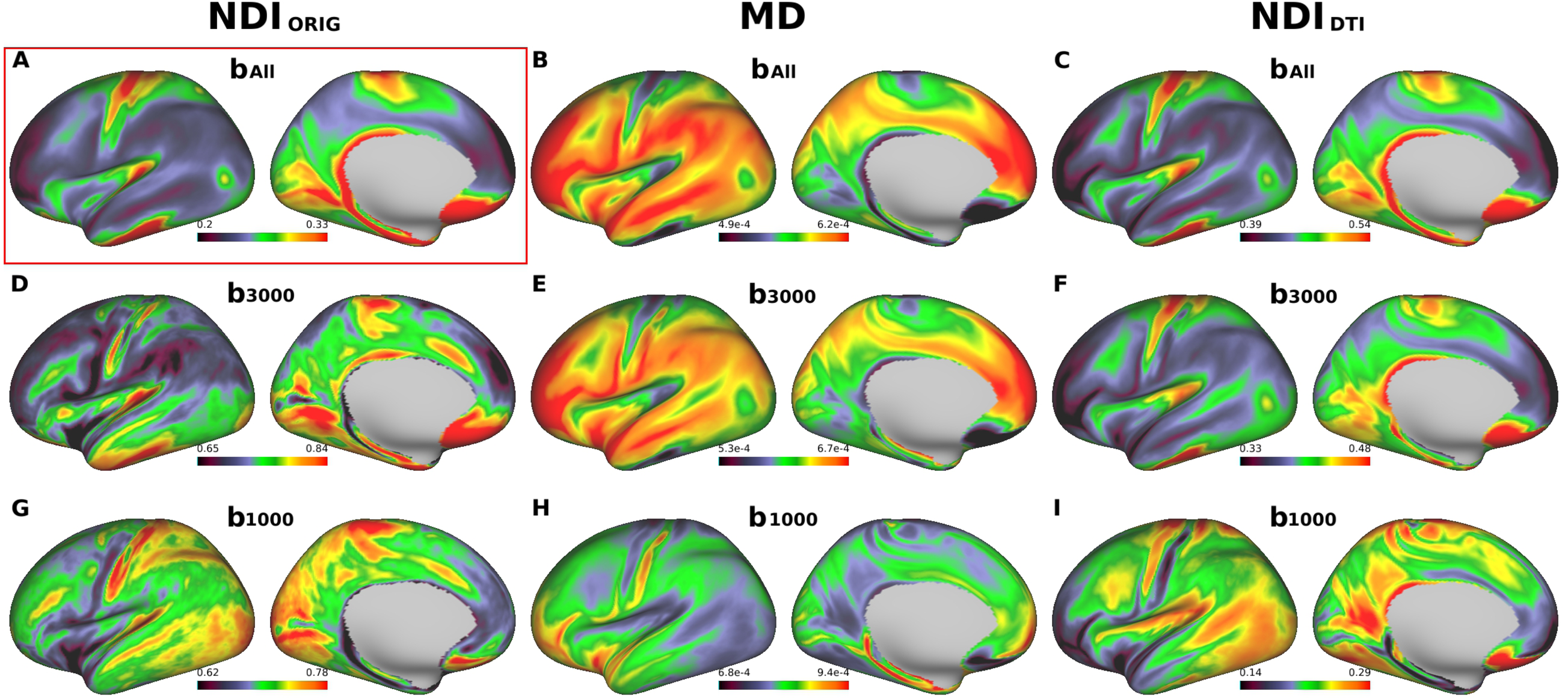
Cross-subject average cortical surface maps of neurite density index (NDI) and mean diffusivity (MD). Cortical surfaces are different in terms of computation methods: original NODDI NDI (NDIORIG) (A, D, G), DTI-derived MD (B, E, H) and DTI-derived NODDI NDI (NDIDTI) (C, F, I) with different b-shell datasets used: all three b-values (bAll), only those of b=3000 (b3000) and b=1000 (b1000), respectively. Reference cortical surface maps of NDIORIG with bAll in (A) showed high values in primary sensorimotor, visual, auditory cortices as well as the middle temporal (MT) area, similar to the cortical myelin distribution as reported previously20. Both MD/bAll and MD/b3000 (B, E) showed inversed appearance to the reference, as well as both NDIDTI/bAll and NDIDTI/b3000 (C, F) showed very similar surface distribution to the reference. Note that NDIORIG/b3000 in (D) showed a different pattern from the reference and any computation methods using b1000 (G, H, I) did not show comparable pattern with the reference. https://balsa.wustl.edu/L66BP

**Figure 3.**
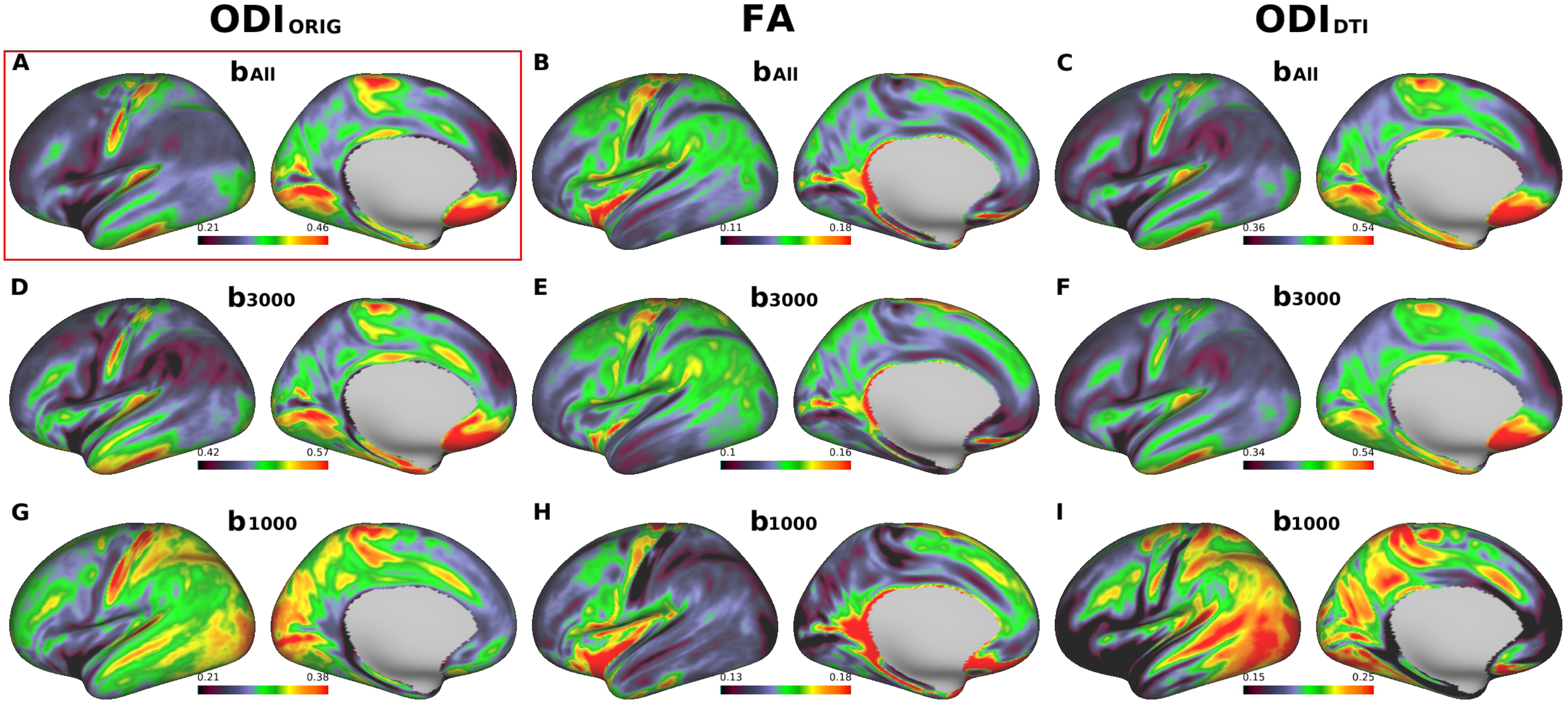
Cross-subject average cortical surface maps of orientation dispersion index (ODI) and fractional anisotropy (FA). Cortical surfaces are different in terms of computation methods: original NODDI ODI (ODIORIG) (A, D, G), DTI-derived FA (B, E, H) and DTI-derived NODDI ODI (ODIDTI) (C, F, I), each used different b-shell datasets: all three b-values (bAll) vs only those of b=3000 (b3000) and low b-values (b1000), respectively. A reference cortical map of ODI (ODIORIG/bAll) in (A) showed high values in the early sensory areas including somatosensory, auditory, and visual. Note that ODIORIG/b3000, ODIDTI/bAll in (C) and ODIDTI/b3000 in (F) showed similar distribution to the reference. Any computation methods using b1000 (H, I, J) did not show comparable pattern with the reference. Data at https://balsa.wustl.edu/pkkKj

When using same three-shell dataset (b_All_), cortical distribution of MD showed extremely inversed appearance to NDI_ORIG_ (Fig. 2B) as reported previously^20^ — this was also true when using high b-value one-shell dataset (b_3000_) (Fig. 2E), but not when using low b-value one-shell dataset (b_1000_) (Fig. 2H). In correlation analysis using the parcellated data (see Methods & Materials 2.1.3), MD strongly negatively correlated with NDI_ORIG_ when using three-shell dataset (b_All_) (R=−0.96, p<0.00001) and high b-value one shell (b_3000_) (R=−0.86, p<0.00001), but did not strongly correlate when using low b-value one-shell dataset (b_1000_) (R=−0.31, p<0.00001) (Fig. 4). As for FA, cortical maps of FA showed moderate inversed appearance to ODI_ORIG_ and (negative) correlation with ODI_ORIG_ among all datasets (R=−0.40 ∼ −0.62) (Fig. 3, 4). There were not strong correlations between FA and NDI_ORIG_ in any b-shell dataset (R=0.15 ∼ 0.28) (Fig. 4).

**Figure 4.**
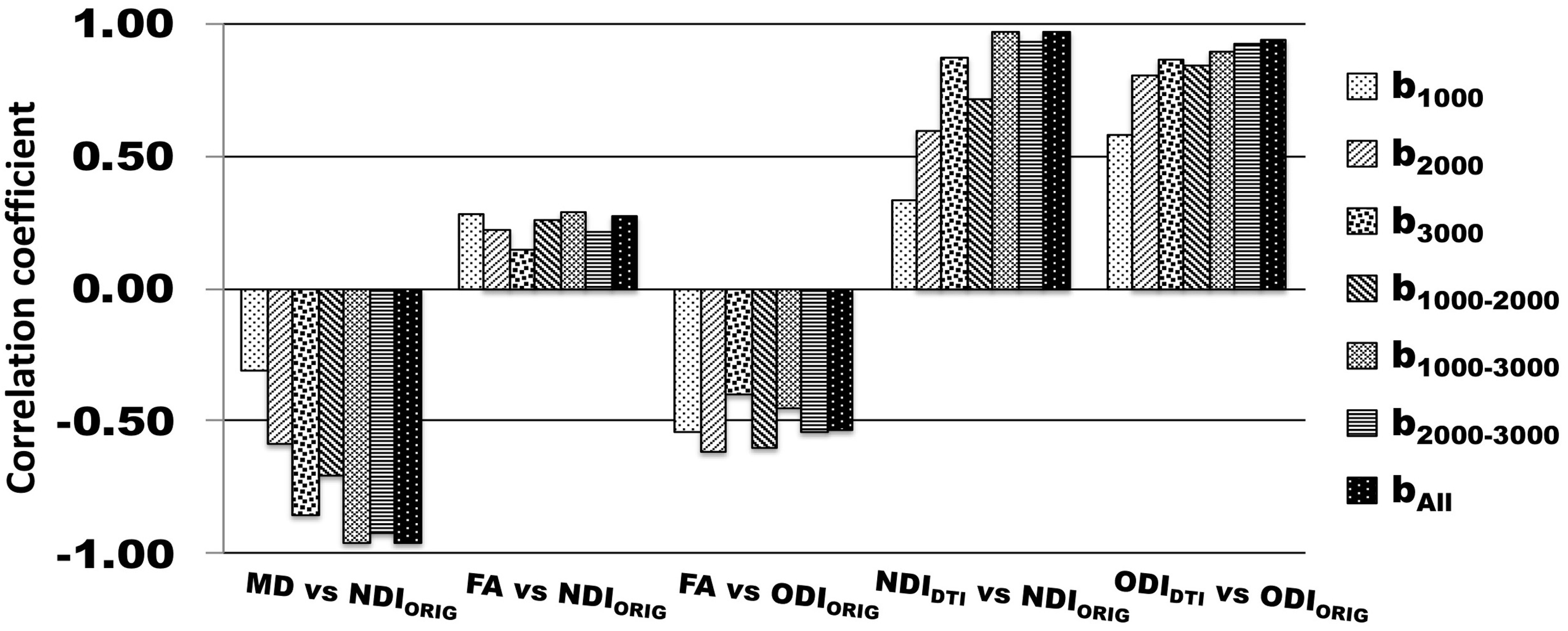
Correlation coefficients of DTI-derived parameters (MD and FA) and DTI-derived NODDI parameters (NDIDTI and ODIDTI) with the references that are the original NODDI parameters on three-shell dataset (NDIORIG/bAll and ODIORIG/bAll). Correlation coefficients to the references were calculated using averaged surface maps across all subjects in each different b-shell dataset type (b1000, b2000, b3000, b1000-2000, b1000-3000, b2000-3000 and bAll). All correlations were significant (p<0.00001).

DTI-derived NODDI maps (NDI_DTI_, ODI_DTI_) also showed very similar cortical distributions of NDI and ODI in average surface maps across all subjects (Fig. 2A, C, F for NDI_DTI_ and Fig. 3A, C, F for ODI_DTI_), particularly when using high b-value dataset including b_All_ and b_3000_. The correlation analysis showed that correlation coefficients between the DTI-derived NODDI and original NODDI parameters were extremely high for both NDI (NDI_DTI_/b_All_: R=0.97, NDI_DTI_/b_3000_: R=0.87, p<0.00001) and ODI (ODI_DTI_/b_All_ R=0.94, ODI_DTI_/b_3000_ R=0.86, p<0.00001) (Fig. 4).

To investigate the agreement of DTI-derived NODDI compared with the original NODDI, the Bland-Altman analysis was applied to the values of cortical parcellations using those of complete data and original NODDI as a reference. When all of the dMRI data (b_All_) were used, the results of DTI-derived NODDI showed a consistent bias: NDI_DTI_ overestimated by a difference of around 0.20 and ODI_DTI_ by 0.15 to 0.10 as compared with those of original NODDI (Fig. 5 A, C). The NDI_DTI_/b_3000,_ (Fig. 5B) also showed a consistent bias, which was a little smaller than NDI_DTI_/b_All_ (Fig. 5A). The bias of ODI_DTI_/b_3000_ (Fig. 5D) was almost same as in the three-shell dataset (Fig. 5C).

**Figure 5.**
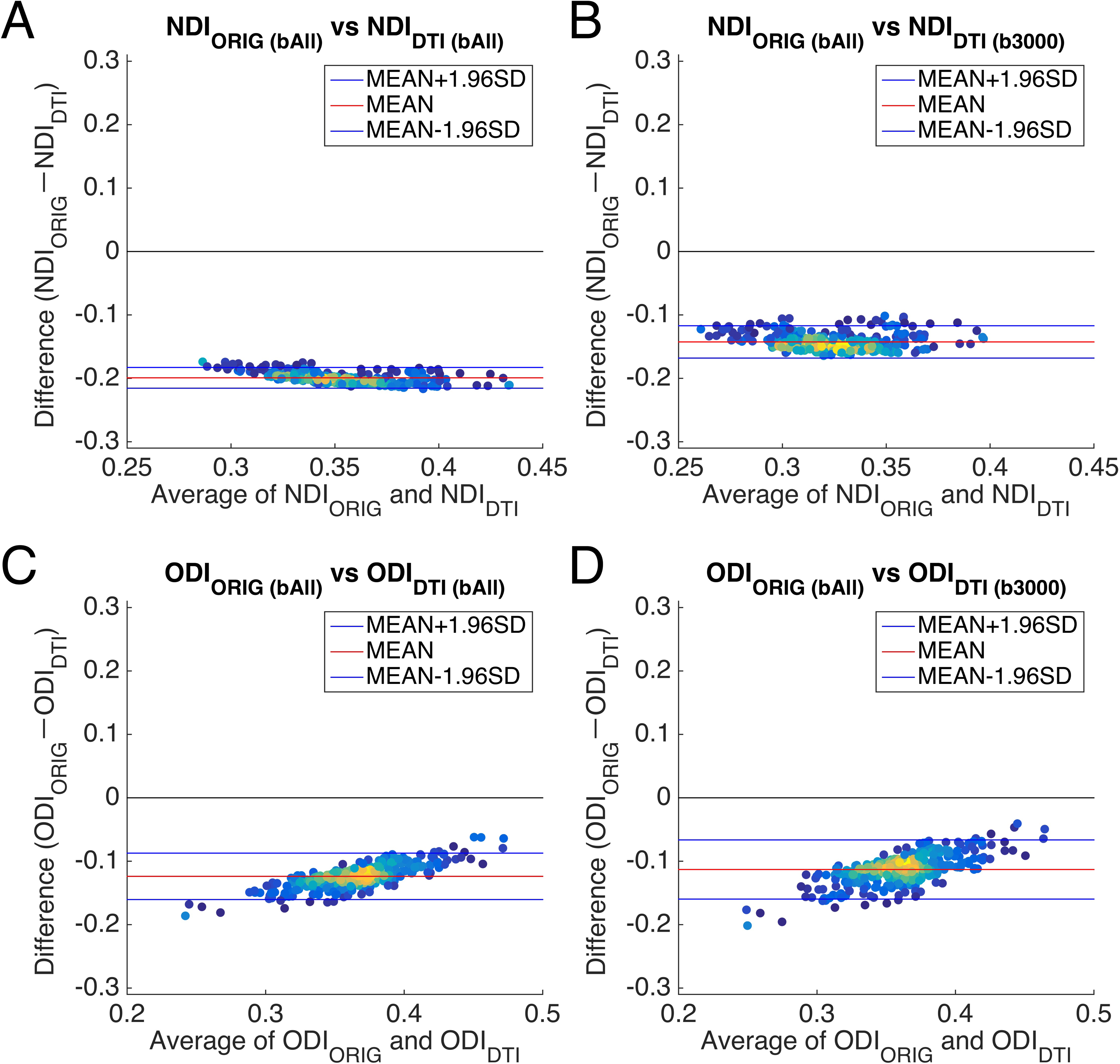
Bland-Altman plots between DTI-derived NODDI and original NODDI parameters in vivo. (A and C) show Bland-Altman plots between DTI-derived NODDI parameters in the three-shell dataset (bAll) and the original NODDI parameters in the three-shell dataset (bAll). (B and D) show Bland-Altman plots between DTI-derived NODDI parameters in the high b-value one-shell dataset (b3000) and the original NODDI parameters in the three-shell dataset (bAll). Plots are coloured by their density. Blue lines show the mean±1.96*SD and the red line shows the mean value. Abbreviations; NDIORIG: neurite density index estimated using the original NODDI model, ODIORIG: orientation dispersion index estimated using the original NODDI model, NDIDTI: neurite density index estimated using DTI-derived NODDI, ODIDTI: orientation dispersion index estimated using DTI-derived NODDI.

Other datasets including a high b-value shell also provided comparable results with the original NODDI (Fig. S3-4). If b=3000 is included in the two-shell data (b_1000-3000_ and b_2000-3000_), both NDI_DTI_ and ODI_DTI_ showed a similar surface distribution to the reference (Fig. S3 A, D, F, Fig. S4 A, D, F). The correlation coefficients were very high in the group-wise maps for both NDI_DTI_ and ODI_DTI_ (b_1000-3000_: R=0.97, R=0.89, b_2000-3000_: R=0.93, R=0.92, respectively, p<0.00001) (Fig. 4). The Bland-Altman analysis showed that the dataset of high and low b-value two-shell (b_1000-3000_) (Fig. S5) had a constant bias of NDI_DTI_ and slightly upward sloping bias of ODI_DTI_, which were almost the same size as in the three-shell dataset. High b-value two-shell (b_2000-3000_) (Fig. S5 A) had also a constant bias of NDI_DTI_ but with a somewhat smaller size than that in three-shell dataset (b_All_). The bias of ODI_DTI_ was almost same size as in the three-shell dataset (Fig. 5 C, S5 B).

The dataset not including a high b-value shell showed inconsistent cortical distributions with the reference. For the two-shell dataset (b_1000-2000_), NDI_DTI_ was a little different and the correlation coefficient was moderate (R=0.71, p<0.00001) (Fig. 4), while ODI_DTI_ showed relatively high correlations in the group-wise maps (R=0.84, p<0.00001) (Fig. 4). One-shell datasets using lower b-value shells (i.e. b_1000_ and b_2000_) did not provide comparable surface maps of NDI_DTI_ (Fig. S3 L, N) and ODI_DTI_ (Fig. S4 L, N). For example, for the low b-value one-shell dataset (b_1000_), both NDI_DTI_ and ODI_DTI_ showed different surface distributions from the reference (Fig. S3 A, N, Fig. S4 A, N), as well as very low correlation coefficients for NDI_DTI_ (R=0.33 p<0.00001) and ODI_DTI_ (R=0.58, p<0.00001) (Fig. 4). This trend was also found when using the middle high b-value one-shell dataset (b_2000_). Only ODI_DTI_ showed a similar surface distribution to the reference (Fig. S4 A, L) and high correlation coefficients (R=0.80, p<0.00001) (Fig. 4), while NDI_DTI_ showed different surface distribution from the reference (Fig. S3 A, L) and relatively low correlations (R=0.59, p<0.00001) (Fig. 4).

When comparing original NODDI parameters using one- or two-shell datasets to the reference, any two-shell datasets provided similar surface maps of NDI_ORIG_ and ODI_ORIG_ with the reference and they were highly correlated (R>0.91, p<0.00001) (Fig. S6). However, NDI_ORIG_ using low b-value one-shell datasets were not significantly correlated to the reference (R<0.19, p>0.00001), while ODI_ORIG_ was relatively correlated even though using one-shell datasets (R>0.71, p<0.00001) (Fig. S6), as show in the simulation study in Zhang et al^5^.

### 3.2. Simulation for the effect of heterogeneity in CSF volume fraction on parameters of NODDI, DTI and DTI-derived NODDI

We simulated the percent changes in NODDI, DTI and DTI-derived NODDI depending on altered CSF compartment (V_iso_) and b-shell dataset. As compared with the reference condition (V_iso_=0.1), apparent differences in amount of change in the parameters were found across type of calculation (NODDI, DTI, DTI-derived NODDI) and b-shell schemes (Fig. 6). While the original NODDI using all the b-shell dataset (b_All_) was reasonably unbiased by altered V_iso_, the parameters of DTI and DTI-derived NODDI tended to be largely biased particularly used b-shell datasets not including high b-value volumes (b=3000) (b_1000_, b_2000_, and b_1000-2000_) (Fig. 6). The dataset including high b-value (b_3000_, b_1000-3000_, b_2000-3000_ and b_All_) were relatively less biased across ranges of V_iso_ changes. These findings suggest that the error of DTI derived parameters is sensitive to the heterogeneity of CSF partial voluming, particularly when lower b-value data was applied. To confirm the specificity of this findings, we also performed another simulation in which ‘homogeneous’ but small CSF volume fraction was assumed (Supplementary Text 3.2). This confirmed that correlation between original NODDI and DTI-derived NODDI was reasonably high as long as CSF volume fraction is not ‘heterogeneous’.

**Figure 6.**
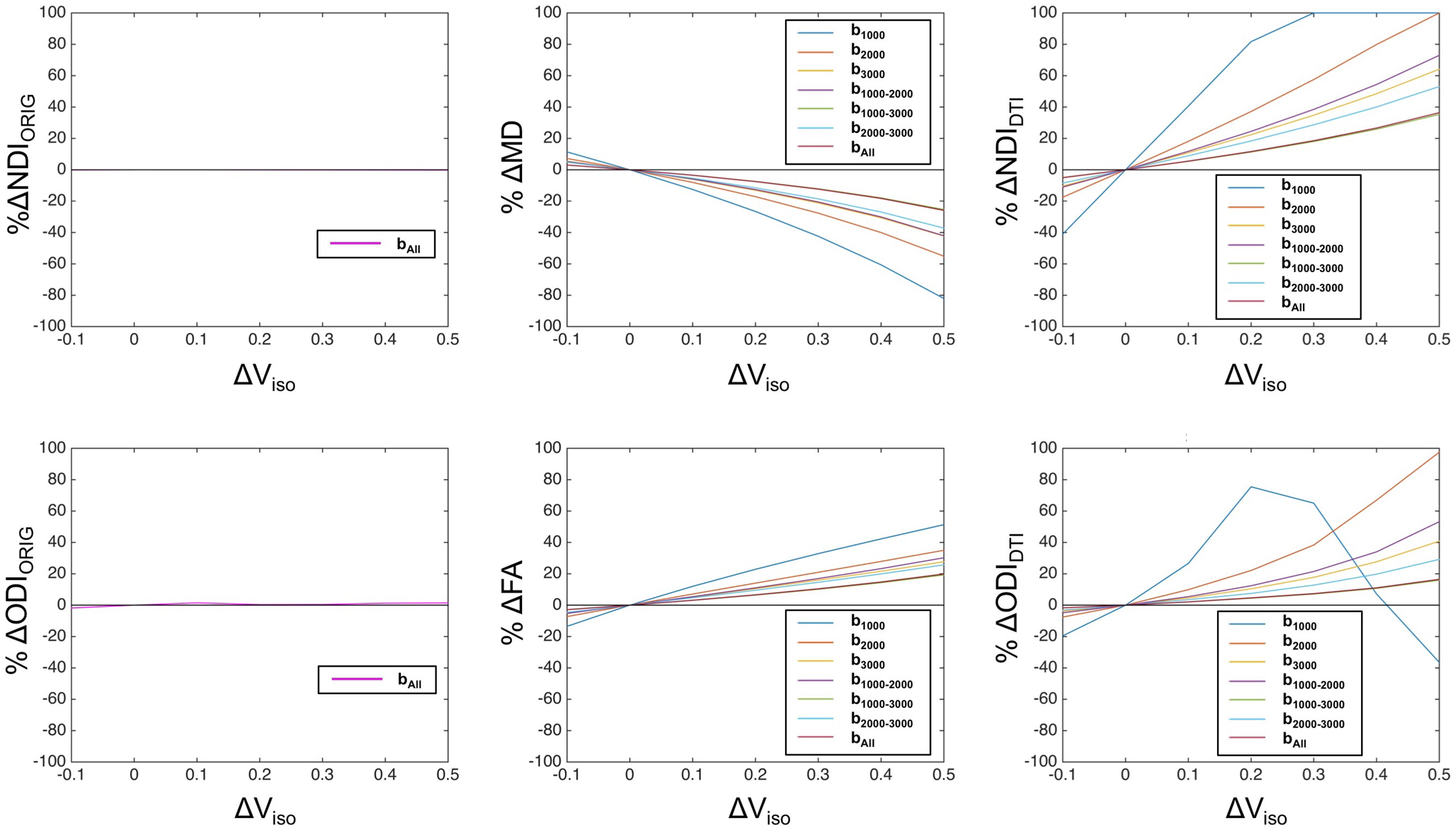
Results of simulation for percent errors in DTI-derived parameters depending on various range of the CSF volume fraction. A) The percent errors in NDI in NODDI (%ΔNDIORIG, left panel), DTI MD (%ΔMD, middle panel) and DTI-derived NDI (%ΔNDIDTI, right panel) against various levels of CSF volume fraction (Viso) relative to the refence value (=0.1). B) The percent errors in ODI from the NODDI (%ΔODIORIG, left panel), DTI FA (%ΔFA, middle) and DTI-derived ODI (%ΔODIDTI, right panel). Dataset types of b-shell schemes b1000, b2000, b3000, b1000-2000, b1000-3000, b2000-3000 and bAll are shown in different colored lines as in the legend in each panel. Note that the one-shell low b-value data set (b1000) shows the largest size of errors in DTI and DTI-derived NODDI parameters among all the datasets, which suggests high sensitivity to partial volume effects in the cortical gray matter. The smallest change in DTI-derived NODDI and DTI parameters was found when using the three-shell dMRI data (bAll), followed by the high b-value two-shell (b1000-3000) and one-shell dMRI data (b3000). Abbreviations; NDIORIG: original NODDI neurite density index, ODIORIG: original NODDI orientation dispersion index, NDIDTI: DTI-derived NODDI neurite density index, ODIDTI: DTI-derived NODDI orientation dispersion index

## Discussion

Accumulating evidence have suggested that high b-value dMRI signal is more sensitive to neurites and neural tissue changes than low b-value dMRI signal. The in vitro study of optic nerve with q-space analysis of diffusion-weighted spectroscopy^31^ identified slower-decaying component of dMRI signals which have 1) diffusion displacement restricted to ∼2 µm close to axonal diameter, 2) longer T2 than rapid diffusing components including myelin water and 3) dependency on neurite orientation, thus suggesting that this component originated from intra-neurite water. The T2 time for myelin water is very short (10-20ms), while intra- and extra-cellular water is longer than 60ms^46^, thus making dMRI signals with TE=89.5ms insensitive to myelin water. The dependency of high b-value diffusion-weighted signal on neurite orientation is largely caused by neurite membrane rather than by other longitudinal structures including myelin and neurofilaments^47^. Simulation showed that the majority of dMRI signals with b-value less than 1000 s/m^-2^ represents fast-diffusion components which may originate free water such as in CSF compartment. On the other hand, recent high b-value DTI in clinical studies showed higher sensitivity to neurobiological changes than low b-value — apparent diffusion coefficient from high b-value DTI was sensitive to the contrast between gray/white matter^32^, ischemic/hypoxic changes in the gray matter^33^, white matter disintegrity in schizophrenia^34^ and maturation in juveniles^35^. Taken together, signals obtained at high b-value are likely sensitive to neurobiological changes including neurites, however, there is not completely established model that quantitates a specific property of neurites. The DTI and NODDI are among the most widely used models: the former is a simplified linear model that accounts for Gaussian process of diffusion motion of water molecule in the tissue, and the latter explicitly models neurite properties in the tissue based on non-linear nature in high b-value data.

The NODDI is among the most validated models for its predictability of neurite properties. Accumulated histological evidence indicates that NDI and ODI of brain and spinal cord tissues are reasonably correlated with histology-based neurite density^25^ and orientation dispersion^25–28^, respectively. The NODDI ODI is relatively robust against data quality^5^, and well represented the histology-based neurite orientation dispersion in all the studies^25–28^ including a single shell and multi-shell dMRI. It is notable that by applying multi-shell dMRI in the spinal cord specimen of multiple sclerosis. Grussu et al. ^25^ revealed that NDI of NODDI was fairly correlated with histology-based density of neurites as assessed by staining neurofilaments. However, since the intrinsic diffusivity is simplified^5^ (see section of Limitation), careful attention is needed for potential bias depending on the diffusivity, for example, when mixing analysis across tissue subtypes such as gray and white matter^20^. Therefore, we estimated specific tissue subtype, cortical gray matter and analyzed the relation of DTI metrics, DTI-derived NODDI metrics to the original NODDI.

Here, we showed in healthy subjects that cortical metrics of DTI and DTI-derived NODDI parameters were highly correlated with those of original NODDI, when used a particular set of b-shell scheme in dMRI. 1) the DTI MD was negatively correlated with NODDI NDI when data included high b-value (b=3000) (R>0.9), 2) the DTI FA was partially correlated with NODDI ODI and NDI particularly when used middle to lower ranged b-value (b=1000-2000) (R>0.87), 3) both NDI and ODI of DTI-derived NODDI showed high correlation with the original NODDI (R>0.9) when used dMRI data including high b-value (b=3000). Simulation analysis suggests that less relation of DTI to NODDI when used low b-value data is due to higher sensitivity to heterogeneity in CSF volume fraction in the intra-tissue and/or partial volume. The HCP data and simulation showed that high b-value dMRI data resulted in a constant numerical bias, i.e. same amount of error over the range of values, potentially due to the bias in the DTI measures coming from non-Gaussian distribution of high b-value dMRI data. Since high b-value dMRI data is often in non-Gaussian distribution, applying linear DTI model for such high b-value dMRI data may result in biases of calculated measures as compared with those used b=1000 dMRI data. Past literature also notes that when using high b-value dMRI data, the values of MD were underestimated^48^ and those of FA were overestimated^4,49,50^ compared with those of b=1000 dMRI data. We’ve also confirmed in the simulation that the biases are constant over a possible range of values between DTI-derived NODDI and original NODDI (data not shown). Therefore, the bias of DTI may be due to the effect of kurtosis of high b-value dMRI data. These findings indicate that DTI parameters in cortical gray matter are highly related to those of NODDI when analyzed using high b-value dMRI data and are not very predictive when used low b-value dMRI. This suggests that analyzing cortical microarchitecture by both DTI and NODDI is redundant and does not surpass usefulness of cortical neurite mapping by either way.

The DTI and DTI-derived NODDI were sensitive to the errors caused by heterogeneity of CSF volume fraction and b-shell scheme of the data. When not using the high b-value shell, the cortical distribution of DTI-derived NODDI parameters showed completely different pattern from those of original NODDI (Fig. S3-4). Our simulation suggests this is because low b-value DTI-derived NODDI parameters are more sensitive to change in V_iso_ due to heterogeneity and partial voluming of CSF (Fig. 6). Low b-value dMRI is theoretically sensitive to fluid signals or ‘T2 shine-through’ effect as well as to tissue diffusivity, whereas high b-value dMRI is more specific to tissue diffusivity^32,51^. In addition, the partial volume effects of CSF may vary across cortical regions according to cortical thickness and their heterogeneity within the cortex. The effect is not completely removed even though the partial volume effect is reasonably reduced by surface-based analysis as compared with volume-based analysis (see Supplementary text 2, Fig. S1). Moreover, the model of DTI by itself does not account for multi compartments in the tissue and also suffers from a partial volume effect of CSF and results in fitting error particularly in the cortex^2,52^. In contrast, the NODDI explicitly considers a CSF compartment s is insensitive to the heterogeneity of CSF as shown in the simulation study (Fig 6). The high b-value DTI and DTI-derived NODDI parameters were also biased in a fixed manner (Fig 5 and Supplementary text 3.2), which are likely caused by non-Gaussian distribution^36^. The values of MD were underestimated and those of FA were overestimated (Supplementary Text 3.2) in high b-value datasets, consistent with previous studies for MD^48^ and FA^4,49,50^.

The current study gives insights and interpretations into recent studies which applied NODDI and DTI in the same sample. In particular, NODDI and DTI showed different sensitivity to the neurobiological changes of interest, while there is a potential variety of CSF contamination and data sampling. Grussu et al. studied NODDI and DTI in spinal cord in healthy subjects^53^. They applied DTI to low b-value dMRI and NODDI to multi-shell data and found that NODDI ODI was the most sensitive to the contrast between gray and white matter. Kamagata et al., applied both DTI and NODDI parameters in Parkinson’s disease and controls using multi-shell dMRI data^29^. The DTI was calculated using low b-value data (b=1000) and NODDI with 2-shell of b=1000 and 2000. Interestingly, both NDI and ODI of the NODDI metrics in the cortical gray matter were more sensitive to discriminate patients from controls than those of DTI, which may support higher sensitivity of NODDI than low b-value DTI to the neuropathological changes in this disease. These are in line with our result that DTI-derived NODDI parameters with low to middle b-value data was not strongly correlated with the high b-value NODDI parameters. Mah et al. also studied NODDI and DTI in early adolescent brain and found that NODDI NDI was more sensitive to age-related changes as compared to DTI MD^54^. They also showed that subcortical gray matter structures, which may be less affected by partial voluming CSF than cortex, showed high correlation between MD and NDI (R=0.69-0.88) and between FA and ODI (R=0.70-0.81). In addition, Batalle et al. analyzed cortical metrics of NODDI and DTI in infant brain and results were complicated^55^. They applied relatively low-resolution dMRI (2mm) for small sized brain and found the dissociated pattern of changes in the cortical NDI and MD. As expected, MD and NDI showed inversed pattern across ages but only after gestational age of 38-week. Parallel pattern between MD and NDI was found before age of 38-week, which may be due to errors in partial voluming of CSF due to small sized brain and thin cortex. The partial volume effect of CSF may not be negligible in their results, since DTI data was calculated based on the low b-value dMRI data.

Preclinical studies also showed neurobiological changes by NODDI and DTI. Using Alzheimer’s model of transgenic mice, Colgan et al. performed NODDI and DTI using multi-shell dMRI data and found higher sensitivity of NODDI NDI than low b-value DTI MD to histology-based marker of neurodegeneration, tau immunoreactivity^56^. The in vitro study using spinal cord specimen of multiple sclerosis was scanned with multi-shell dMRI including high b-value. They also showed that MD of DTI was negatively correlated with histology-based neurite density in the same specimen. Our previous study which used multi-shell dMRI data also found a very strong relationship between DTI MD and NODDI NDI and that values of FA was also associated with ODI depending on the MD^20^.

This study provides important implications for future dMRI studies. First, it is redundant to apply both DTI and NODDI to dMRI data for estimating microstructure in cortical gray matter. As formulated by a mathematical conversion from DTI to NODDI, there is no additional quantitative information available by applying both. When used low b-value dataset, DTI suffers from the errors of CSF heterogeneity as compared with NODDI. Therefore, users can choose either method depending on the dMRI data acquired, and neurobiological significance may not be changed. Second, a high b-value DTI is potentially useful for cortical neurite mapping, particularly in clinical setting. A shorter dMRI scan will be particularly helpful for clinical patients such as patients with Alzheimer’s disease who cannot keep still long time. DTI can be estimated with relatively few directions - at least 6 or in general more than 30 are recommended ^57^, whereas the original NODDI is recommended with at least 90 directions ^5^, which means three times higher efficiency. HCP-style scanning with high spatial resolution dMRI with 30 directions does not exceed 3 min. Third, low b-value DTI is not appropriate for cortical mapping and suffer from errors from heterogeneity and partial voluming of CSF. The heterogeneity itself cannot be completely estimated by the currently available resolution of dMRI. A special sequence, such as ‘fluid-attenuated inversed recovery DTI’, can be useful by reducing CSF signals ^58,59^. Meanwhile, the NODDI is more robust against the errors from CSF partial voluming for cortical mapping of microarchitecture.

One of limitations of this study is that it relies on the relative validity of NODDI over DTI. Recently, there is subject of debate about the eligibility on simplification in the NODDI associated with constrained intrinsic diffusivity^24,60^. There is an attempt to develop a novel method that explicitly analyzes local complexities of diffusivity^24^. The method is potentially useful for future application; however, it is technically demanding for scanning, particularly, specific diffusion gradient encoding both in linear tensor and spherical tensor. There is also need for investigations on whether the intrinsic diffusivity is significantly changed in-vivo and how it influences the quantification of neurite properties in the gray matter. In the previous study, we optimized the intrinsic diffusivity for the gray matter (1.1 × 10^-3^ mm^2^/s) based on non-linear multiparametric fitting^20^, which resulted in reasonable findings of correlation between neurite density and myelin contrast as expected from histological evidence^61,62^. Second, we did not analyze the relationship between DTI and NODDI in the ‘white matter’ in this article. We estimated that the volume fraction of isotropic diffusion is larger in the white matter (0.21±0.1) than in the gray matter (0.09±0.06) (see also Supplementary Text 2), as expected from the fact that the white matter is also a major site for convective flow of CSF^63^. Despite potential larger fraction of CSF, the white matter is not affected by partial voluming of CSF in the subarachnoid space like in the cortical gray matter. Therefore, as long as the volume fraction of CSF is relatively heterogenous across regions in the white matter, the relationship of NODDI and DTI may be similar to those in the gray matter. This is also supported by the current simulation studies on heterogeneity of CSF volume fraction as shown in section 3.2 and Supplementary Text 3.2. Third, we applied DTI model for multi-shell dMRI datasets, which are known to have non-Gaussian distribution. Mathematically, non-linear model like DKI^39^ is more suitable for such non-Gaussian dMRI data than DTI. There is also evidence that parameters of DTI such as FA and MD are comparable with those from DKI^64^, however, we did not include the comparison of DKI with NODDI in this study because of limited space.

## 5. Conclusion

For addressing cortical microarchitecture, conventional DTI with low b-value dataset is not very useful because of contamination with the heterogeneity of CSF, whereas NODDI is robust against these factors. Cortical DTI parameters were closely associated with those of NODDI, particularly using data including high b-value data. DTI-derived NODDI based on high b-value dataset showed remarkably similar cortical distributions with those of NODDI, supporting the previous notion of the mathematical conversion between the DTI and NODDI. Simulation also supported these findings that potential intra-tissue CSF fraction and partial voluming of arachnoid CSF may be causing the error and bias in the cortical maps different from those of original NODDI. Although its similarity, analyzing with high b-value dataset and DTI does not add more information for cortical microarchitecture than NODDI.

## 6. Notes

Data of figures and supplementary figures are available at https://balsa.wustl.edu/study/show/G331M

## 7. Ethics Statement

The use of HCP data in this study was approved by the institutional ethical committee (KOBE-IRB-16-24).

## 8. Author Contributions

HF and TH designed the study. HF, KM and TH made programs for analysis. HF, TH, TA, KF, TY, TO, and KT analyzed the data. HF, TH, MG, HZ, JA, and DV contributed to writing the manuscript. All authors have read and approved the final manuscript.

## 10. Acknowledgements

The data of this study were provided by the Human Connectome Project, WU-Minn Consortium (Principal Investigators: David Van Essen and Kamil Ugurbil; 1U54MH091657) funded by the 16 NIH Institutes and Centers that support the NIH Blueprint for Neuroscience Research; and by the McDonnell Center for Systems Neuroscience at Washington University. This research is partially supported by the program for Brain Mapping by Integrated Neurotechnologies for Disease Studies (Brain/MINDS) and Brain/MINDS-beyond from Japan Agency for Medical Research and development, AMED (JP18dm0207001, JP19dm0307006h0002), by RIKEN Compass to Healthy Life Research Complex Program from Japan Science and Technology Agency, JST, by MEXT KAKENHI Grant (16H03300, 16H03306) (T.H.).

