## Supplementary File for "Diffusion Tensor Model links to Neurite Orientation Dispersion and Density Imaging at high b-value in Cerebral Cortical Gray Matter"

[5]Siemens Healthcare K.K. Japan

[6]Centre for Medical Image Computing and Department of Computer Science, University College London, UK

[7]RIKEN Compass to Healthy Life Research Complex Program, Kobe, Japan

Incl.

**Supplementary Text 1-3**

**Supplementary Table S1**

**Supplementary Figures S1-S6**

**References**

**Supplementary Text**

### 1. Models of NODDI and DTI-derived NODDI

#### 1.1 *The original NODDI Model*

The NODDI method models brain microarchitecture in three compartments that have different properties of water molecules' diffusion motion: the intracellular compartment (restricted diffusion bounded by neurites), the extracellular compartment (outside of neurites and potentially including glial cells), and the CSF compartment [1]. The intracellular compartment is modeled as a set of sticks, i.e., cylinders of zero radius in which diffusion of water is highly restricted in directions perpendicular to neurites and unhindered along them [2–4]. The orientation distribution of these sticks is modeled with a Watson distribution, because it is the simplest distribution that can capture the dispersion in orientations [5]. The extracellular compartment is modeled with anisotropic Gaussian diffusion parallel to the main direction. The CSF compartment is modeled as isotropic Gaussian diffusion. The full normalized signal A is thus written as:

$$A= (1-V_{iso})\{V_{ic} A_{ic}+(1-V_{ic})A_{ec}\}+V_{iso} A_{iso}, \qquad (1)$$

where $A_{iso}$ and $V_{iso}$ are the normalized signal and volume fraction of the CSF compartment; the volume fraction of non-CSF compartment ($1-V_{iso}$) is further divided into intracellular compartment ($V_{ic}$) (=NDI) and extracellular compartment ($1-V_{ic}$); $A_{ic}$ and $A_{ec}$ is the normalized signal of the intracellular and extracellular compartments, respectively. Additional NODDI parameters are isotropic diffusivity ($d_{iso}$) and intrinsic free diffusivity ($d_{\parallel}$) that means the diffusivity parallel to neurites and is constrained. Detailed expressions of mathematical equations and derivation are described in the Appendix in Supplementary material, and these formulations were used for the simulation study.

#### 1.2 *The DTI-derived NODDI calculation*

The equations that relate NODDI to DTI models are detailed in previous studies [6,7]. Briefly, the NDI and the orientation parameter ($\tau$) can be expressed by using DTI measures such as MD and FA in the following equations, assuming that the CSF compartment ($V_{iso}$) is negligible:

$$NDI = 1 - \sqrt{\frac{1}{2}\left(\frac{3MD}{d_{//}} - 1\right)} \qquad (2)$$

$$\tau = \frac{1}{3}\left(1 + \frac{4MD \cdot FA}{|d_{//}-MD|\sqrt{3-2FA^2}}\right), \qquad (3)$$

where $d_{//}$ is a constant for intrinsic diffusivity assumed in the NODDI model. The orientation dispersion index (ODI) is calculated using the following formulas:

$$\tau = \frac{1}{\sqrt{\pi\kappa}\exp(-k)erfi(\sqrt{\kappa})} - \frac{1}{2\kappa} \quad (4)$$

$$ODI = \frac{2}{\pi}\arctan\left(\frac{1}{\kappa}\right), \quad (5)$$

where *erfi* is the imaginary error function and *arctan* is the arctangent. Based on these equations, once we have DTI measures such as FA and MD, 1) NDI can be analytically estimated from MD using formula (2) ($NDI_{DTI}$) by using an assumed value of $d_{//}$, 2) $\tau$ can be calculated using formula (3) and values of MD and FA, 3) $\kappa$ can be estimated using formula (4) by using a look-up-table and a value of $\tau$ calculated at the previous step, and 4) $ODI_{DTI}$ was calculated using the formula (5) and $\kappa$.

The values of DTI and predicted NODDI parameters based on Eq. (2)-(5) were plotted in Fig. 1 in the main text using in-house script written by MATLAB (R2013a) (http://www.mathworks.com/). We used $d_{//}=1.1 \times 10^{-3}$ mm$^2$/s (optimized for gray matter [8]) and for an expected range of MD in the cortex (5 to 6 x $10^{-4}$ mm$^2$/s, see Fig. 4B in our previous study[8]). We found that the value of NDI is predicted by a monotonically increasing function of the inverse of MD (Fig. 1A in the main text) and that ODI is a monotonically decreasing function depending both on MD and FA (Fig. 1B in the main text).

### 2. Contamination of CSF in cortical surface-based and volume-based analysis in the NODDI model

Looking at a volume slice of the $V_{iso}$ (isotropic diffusion compartment) of NODDI, it is notable that the cerebral cortical ribbon tends to take lower values than the white matter and extra-brain CSF areas. (Fig. S1 A). However, measurement in cortical gray matter areas can easily suffer from partial volume effects because of the limited spatial resolution of dMRI data and can also be influenced by the methods used to resample the data to the surface or volume. Although we applied the surface mapping method (myelin-style) the least affected by the partial voluming, it may be worth estimating whether resampling methods can affect the CSF values in the cortex. Therefore, we evaluated and compared the distribution of CSF values in the whole cerebral cortex between surface and volume-based methods. In the surface mapping method, the values of $V_{iso}$ in the cerebral cortical surface were reasonably low and distributed in a narrow range (mean=0.096, standard deviation=0.063) (Fig. S1 B), while in volume-based analysis, both were relatively high (mean=0.17, standard deviation=0.15) (Fig. S1 B). These findings suggest that the surface mapping method can better avoid the effects of partial voluming as compared with volume-based analysis when using NODDI in cortical gray matter. We also found the values of $V_{iso}$ in the white matter region was mean=0.21, standard deviation=0.097 (Fig. S1 B).

### 3. Simulation for the error in NODDI-DTI

#### *3.1. Formulation and derivation of NODDI model for simulation study*

In this section, we described formulation and derivation of the NODDI model, by which simulation study was performed. In the NODDI model, the signal (*A*) of the tissue is composed of CSF ($A_{iso}$), extracellular ($A_{ec}$) and intracellular compartments ($A_{ic}$) [1] as in Eq. 1. The signal is also dependent on volume fractions of the CSF compartment ($\nu_{iso}$.) and the intracellular compartments ($\nu_{ic}$). We describe in detail how each of $A_{iso}$, $A_{ec}$, and $A_{ic}$ can be expressed mathematically. We also describe how the Watson distribution can be expressed by a mathematical equation.

##### *3.1.1. CSF compartment ($A_{iso}$)*

Since $A_{iso}$ is dependent on isotropic diffusion, it can be expressed as

$$A_{iso} = e^{-bd_{iso}}, \qquad (A1)$$

where $b$ is b-value of dMRI and $d_{iso}$ is the diffusion coefficient of the CSF.

##### *3.1.2. Extracellular compartment ($A_{ec}$)*

According to Zhang et al. [1], $A_{ec}$ is expressed as follows:

$$A_{ec} = \exp\left(-b\boldsymbol{q}^T \cdot \int_{\mathbb{S}^2} f(\boldsymbol{n}|\boldsymbol{\mu},\kappa)D(\boldsymbol{n})\, d\boldsymbol{n} \cdot \boldsymbol{q}\right), \qquad (A2)$$

where $\boldsymbol{q}$ is an unit vector which is the direction of diffusion weighting gradient and $D(\boldsymbol{n})$ is a cylindrical symmetry tensor whose main axis is along the direction of $\boldsymbol{n}$.

On the other hand, according to Zhang et al. [1], let $d_{\parallel}$ and $d_{\perp}$ be the diffusion coefficients which are parallel and perpendicular to the main axis in the intracellular compartment, respectively. The diffusion coefficients ($d'_{\parallel}$ and $d'_{\perp}$) which are parallel and perpendicular to the main axis in the extracellular compartment, are expressed as follows:

$$\begin{cases} d'_{\parallel} = d_{\parallel} - d_{\parallel}\nu_{ic}(1-\tau_1) \\ d'_{\perp} = d_{\parallel} - d_{\parallel}\nu_{ic}\left(\dfrac{1+\tau_1}{2}\right), \end{cases} \qquad (A3)$$

where $\tau_1$ is expressed as follows [1]:

$$\tau_1 = -\frac{1}{2\kappa} + \frac{1}{\sqrt{\pi\kappa} \cdot e^{-\kappa} \cdot erfi(\sqrt{\kappa})}, \quad (A4)$$

where $erfi(x)$ is the incomplete error function and given as below:

$$erfi(x) = \frac{2}{\sqrt{\pi}} \int_0^x e^{t^2} dt. \quad (A5)$$

Since the principal axis of the extracellular compartment is assumed to be parallel to the z-axis, $\boldsymbol{D_{ec}}(\hat{\boldsymbol{z}}, \kappa)$ is expressed as below:

$$\boldsymbol{D_{ec}}(\hat{\boldsymbol{z}}, \kappa) = \begin{pmatrix} d'_{\perp} & 0 & 0 \\ 0 & d'_{\perp} & 0 \\ 0 & 0 & d'_{\parallel} \end{pmatrix}. \quad (A6)$$

Therefore, $A_{ec}$ is rewritten using Eq. (A2), (A6) as below:

$$A_{ec} = exp(-b\boldsymbol{q}^T \cdot \boldsymbol{D_{ec}}(\boldsymbol{\mu}, \kappa) \cdot \boldsymbol{q}). \quad (A7)$$

Since $\boldsymbol{D_{ec}}(\boldsymbol{\mu}, \kappa)$ is a cylindrically symmetric tensor whose principal axis is in the direction of the principal axis of the Watson distribution (described in detail Appendix 4), namely $\boldsymbol{\mu}$, $\boldsymbol{q}^T \cdot \boldsymbol{D_{ec}}(\boldsymbol{\mu}, \kappa)\boldsymbol{q}$ is a function of $\theta = \boldsymbol{q} \cdot \boldsymbol{\mu}$ which is the relative angle between the principal axes of MPG and Watson distribution. Hence, without loss of generality, let $\boldsymbol{\mu} = \hat{\boldsymbol{z}}$. Since $\boldsymbol{D_{ec}}(\hat{\boldsymbol{z}}, \kappa)$ is cylindrically symmetrical to the z-axis in this case, $A_{ec}$ depends only on $\theta = \boldsymbol{q} \cdot \boldsymbol{\mu}$, which is the angle between MPG and z-axis, not on the azimuthal angle $\phi$. Hence, without loss of generality, let $\phi = 0$. Now, let $\boldsymbol{R}(-\theta_q)$ be the rotation matrix, which makes the direction of MPG ($\boldsymbol{q}$) parallel to z-axis,

$$\begin{aligned} \boldsymbol{q}^T \cdot \boldsymbol{D_{ec}}(\hat{\boldsymbol{z}}, \kappa)\boldsymbol{q} &= \left(\boldsymbol{R}(-\theta_q) \cdot \boldsymbol{q}\right)^T \cdot \boldsymbol{D_{ec}}\left(\boldsymbol{R}(-\theta_q) \cdot \hat{\boldsymbol{z}}, \kappa\right) \cdot \left(\boldsymbol{R}(-\theta_q) \cdot \boldsymbol{q}\right) \\ &= \hat{\boldsymbol{z}}^T \cdot \boldsymbol{D_{ec}}\left(\boldsymbol{R}(-\theta_q) \cdot \hat{\boldsymbol{z}}, \kappa\right) \cdot \hat{\boldsymbol{z}} \\ &= \begin{pmatrix} 0 & 0 & 1 \end{pmatrix} \begin{pmatrix} 1 & 0 & 0 \\ 0 & cos\theta & -sin\theta \\ 0 & sin\theta & cos\theta \end{pmatrix}^T \begin{pmatrix} d'_{\perp} & 0 & 0 \\ 0 & d'_{\perp} & 0 \\ 0 & 0 & d'_{\parallel} \end{pmatrix} \begin{pmatrix} 1 & 0 & 0 \\ 0 & cos\theta & -sin\theta \\ 0 & sin\theta & cos\theta \end{pmatrix} \begin{pmatrix} 0 \\ 0 \\ 1 \end{pmatrix} \\ &= d'_{\perp}sin^2\theta + d'_{\perp}cos^2\theta. \end{aligned} \quad (A8)$$

Summarizing the above, $A_{ec}$ is denoted using Eq. (A7), (A8) as below:

$$A_{ec} = exp\left(-b(d'_{\perp}sin^2\theta + d'_{\perp}cos^2\theta)\right), \quad (A9)$$

where $\theta = \boldsymbol{q} \cdot \boldsymbol{\mu}$.

*3.1.3. Intracellular compartment ($A_{ic}$)*

According to Zhang et al.,

$$A_{ic} = \int_{\mathbb{S}^2} f(\boldsymbol{n}|\boldsymbol{\mu},\kappa) e^{-bd_{\parallel}(\boldsymbol{q}\cdot\boldsymbol{n})^2}\, d\boldsymbol{n}\,, \tag{A10}$$

where $d_{\parallel}$ is intrinsic diffusivity. $A_{ic}$ cannot be expressed by elementary functions. First, the Watson distribution is expanded using spherical harmonics. Let $f_{l0}^{c}(\kappa)$ be an expansion coefficient, when $f(\boldsymbol{n}|\hat{\boldsymbol{z}},\kappa)$ is expanded using spherical harmonics.

$$f(\boldsymbol{n}|\hat{\boldsymbol{z}},\kappa) = \sum_{l=0}^{\infty} f_{l0}^{c}(\kappa) Y_{l0}(\theta_{\boldsymbol{n}},0). \tag{A11}$$

The Watson distribution $f(\boldsymbol{n}|\boldsymbol{\mu},\kappa)$, whose mean orientation is $\boldsymbol{\mu}$, is expressed by using Wigner Rotation Matrix [9] as follows:

$$\begin{aligned}
f(\boldsymbol{n}|\boldsymbol{\mu},\kappa) &= f\left(\boldsymbol{n}|\mathbf{R}\left(-\theta_{\mathbf{q}}\right)\hat{\boldsymbol{z}},\kappa\right) \\
&= \mathbf{R}\left(\theta_{\mathbf{q}}\right) f(\boldsymbol{n}|\hat{\boldsymbol{z}},\kappa) \\
&= \mathbf{R}(\theta_{\mathbf{q}}) \sum_{l=0}^{\infty} f_{l0}^{c}(\kappa) Y_{l0}(\theta_{\boldsymbol{n}},0) \\
&= \sum_{l=0}^{\infty} f_{l0}^{c}(\kappa) \mathbf{R}(\theta_{\mathbf{q}}) Y_{l0}(\theta_{\boldsymbol{n}},0) \\
&= \sum_{l=0}^{\infty} f_{l0}^{c}(\kappa) \sum_{m'=-l}^{l} Y_{lm'}(\theta_{\boldsymbol{n}},\phi_{\boldsymbol{n}}) \sqrt{\frac{4\pi}{2l+1}} Y_{lm'}^{*}(\theta_{\boldsymbol{q}},0),
\end{aligned} \tag{A12}$$

where $\theta_{\boldsymbol{q}}$ is the angle between MPG direction and z-axis.

We substitute this into the $A_{ic}$ (at this time $\mathbf{q} = \hat{\mathbf{z}}$), because if $m \neq 0$, $\int_0^{2\pi} e^{im\phi}\, d\phi = 0$, and if $m = 0, \int_0^{2\pi} 1\, d\phi = 2\pi$.

$$\begin{aligned}
A_{ic} &= \int_{\mathbb{S}^2} \sum_{l=0}^{\infty} f_{l0}^{c}(\kappa) \sum_{m'=-l}^{l} Y_{lm'}(\theta_{\boldsymbol{n}},\phi_{\boldsymbol{n}}) \sqrt{\frac{4\pi}{2l+1}} Y_{lm'}^{*}(\theta_{\boldsymbol{q}},0) e^{-bd_{\parallel}(\hat{\boldsymbol{z}}\cdot\boldsymbol{n})^2}\, d\boldsymbol{n} \\
&= \int_{\mathbb{S}^2} \sum_{l=0}^{\infty} f_{l0}^{c}(\kappa) \sum_{m'=-l}^{l} Y_{lm'}(\theta_{\boldsymbol{n}},\phi_{\boldsymbol{n}}) \sqrt{\frac{4\pi}{2l+1}} Y_{lm'}^{*}(\theta_{\boldsymbol{n}},0) e^{-bd_{\parallel}(\hat{\boldsymbol{z}}\cdot\boldsymbol{n})^2}\, d\boldsymbol{n} \\
&= \sum_{l=0}^{\infty} f_{l0}^{c}(\kappa) \sum_{m'=-l}^{l} \sqrt{\frac{4\pi}{2l+1}} Y_{lm'}^{*}(\theta_{\boldsymbol{q}},0) \int_0^{2\pi} sin\theta_{\boldsymbol{n}} d\phi_{\boldsymbol{n}} \int_0^{\pi} d\theta_{\boldsymbol{n}}\, Y_{lm'}(\theta_{\boldsymbol{n}},\phi_{\boldsymbol{n}}) e^{-bd_{\parallel}cos^2\theta_n}
\end{aligned}$$

$$= \sum_{l=0}^{\infty} f_{l0}^{c}(\kappa) \sum_{m'=-l}^{l} \sqrt{\frac{4\pi}{2l+1}} Y_{lm'}^{*}(\theta_{q}, 0) \int_{0}^{2\pi} sin\theta_{n} d\phi_{n} \int_{0}^{\pi} d\theta_{n} \sqrt{\frac{2l+1}{4\pi}\frac{(l-m')!}{(l+m')!}} P_{l}^{m'}(cos\theta_{n}) e^{im'\phi_{n}} e^{-bd_{\parallel}cos^{2}\theta_{n}}$$

$$= \sum_{l=0}^{\infty} f_{l0}^{c}(\kappa) \sqrt{\frac{4\pi}{2l+1}} Y_{l0}^{*}(\theta_{q}, 0) 2\pi \int_{-1}^{1} dx \sqrt{\frac{2l+1}{4\pi}} P_{l}(x) e^{-bd_{\parallel}x^{2}}$$

$$= 2\pi \sum_{l=0}^{\infty} f_{l0}^{c}(\kappa) \sqrt{\frac{2l+1}{4\pi}} P_{l}(cos\theta_{q}) \int_{-1}^{1} dx\, P_{l}(x) e^{-bd_{\parallel}x^{2}} \qquad (A13)$$

On the other hand, $f_{l0}^{c}(\kappa)$ are expansion coefficients, when $f(\boldsymbol{n}|\hat{\boldsymbol{z}}, \kappa)$ is expressed using spherical harmonics.

$$f(\boldsymbol{n}|\hat{\boldsymbol{z}}, \kappa) = \sum_{l=0}^{\infty} f_{l0}^{c}(\kappa) Y_{l0}(\theta_{n}, 0). \qquad (A14)$$

$f_{l0}^{c}(\kappa)$ can be determined by multiplying $Y_{l'0}^{*}(\theta_{n}, 0)$ and integrating both sides, because of the standard orthogonality of the spherical harmonics.

$$f_{l0}^{c}(\kappa) = \int Y_{l0}^{*} f(\boldsymbol{n}|\hat{\boldsymbol{z}}, \kappa) d\boldsymbol{n}$$

$$= \int \sqrt{\frac{2l+1}{4\pi}} P_{l}(cos\theta) \frac{1}{4\pi} \frac{e^{\kappa(\boldsymbol{\mu}\cdot\boldsymbol{n})}}{M\left(\frac{1}{2}, \frac{3}{2}, \kappa\right)} d\boldsymbol{n}$$

$$= \frac{1}{4\pi} \frac{1}{M\left(\frac{1}{2}, \frac{3}{2}, \kappa\right)} \int_{0}^{2\pi} sin\theta_{n} d\phi_{n} \int_{0}^{\pi} d\theta_{n} \sqrt{\frac{2l+1}{4\pi}} P_{l}(cos\theta_{n}) e^{\kappa cos^{2}\theta_{n}}$$

$$= \frac{\sqrt{2l+1}}{4\sqrt{\pi} \cdot M\left(\frac{1}{2}, \frac{3}{2}, \kappa\right)} \int_{-1}^{1} dx\, P_{l}(x) e^{\kappa x^{2}} \qquad (A15)$$

Now, according to Arfken et al. [10],

$$\int_{-1}^{1} P_{l}(\mu)\, e^{x\mu^{2}} = (x)^{l/2} \frac{\Gamma\left(\frac{l+1}{2}\right)}{\Gamma\left(\frac{2l+3}{2}\right)} M\left(\frac{l+1}{2}, \frac{2l+3}{2}, x\right). \qquad (A16)$$

Hence, $f_{l0}^{c}(\kappa)$ is expressed using Eq. (A15), (A16) as below:

$$f_{l0}^{c}(\kappa) = \frac{\sqrt{2l+1}}{4\sqrt{\pi}} \frac{\Gamma\left(\frac{l+1}{2}\right)}{\Gamma\left(\frac{2l+3}{2}\right)} \frac{M\left(\frac{l+1}{2}, \frac{2l+3}{2}, \kappa\right)}{M\left(\frac{1}{2}, \frac{3}{2}, \kappa\right)} (\kappa)^{l/2}. \qquad (A17)$$

In addition, it can be also applied for factors below, which $A_{ic}$ contains:

$$\int_{-1}^{1} dx\, P_l(x) e^{-bd_{\parallel}x^2} = (-bd_{\parallel})^{l/2} \frac{\Gamma\left(\frac{l+1}{2}\right)}{\Gamma\left(\frac{2l+3}{2}\right)} M\left(\frac{l+1}{2}, \frac{2l+3}{2}, -bd_{\parallel}\right). \tag{A18}$$

In summary, $A_{ic}$ is expressed using Eq. (A13), (A17), (A18) as follows:

$$A_{ic} = 2\pi \sum_{l=0}^{\infty} \frac{\sqrt{2l+1}}{4\sqrt{\pi}} \frac{\Gamma\left(\frac{l+1}{2}\right)}{\Gamma\left(\frac{2l+3}{2}\right)} \frac{M\left(\frac{l+1}{2}, \frac{2l+3}{2}, \kappa\right)}{M\left(\frac{1}{2}, \frac{3}{2}, \kappa\right)} (\kappa)^{l/2} \sqrt{\frac{2l+1}{4\pi}} P_l(cos\theta_q)(-bd_{\parallel})^{l/2} \frac{\Gamma\left(\frac{l+1}{2}\right)}{\Gamma\left(\frac{2l+3}{2}\right)} M\left(\frac{l+1}{2}, \frac{2l+3}{2}, -bd_{\parallel}\right)$$

$$= \frac{1}{4\cdot M\left(\frac{1}{2}, \frac{3}{2}, \kappa\right)} \sum_{l=0}^{\infty} (2l+1) \left( \frac{\Gamma\left(\frac{l+1}{2}\right)}{\Gamma\left(\frac{2l+3}{2}\right)} \right)^2 P_l(cos\theta_q)(-bd_{\parallel}\kappa)^{l/2} M\left(\frac{l+1}{2}, \frac{2l+3}{2}, \kappa\right) M\left(\frac{l+1}{2}, \frac{2l+3}{2}, -bd_{\parallel}\right). \tag{A19}$$

Moreover, the sum of $l$ should be performed for only the even numbers, because the symmetry of θ direction of the Watson distribution.

#### *3.1.4. The Watson distribution*

According to the original NODDI model [1], the Watson distribution is expressed as follows:

$$f(\boldsymbol{n}) = \frac{1}{M\left(\frac{1}{2}, \frac{3}{2}, \kappa\right)} e^{\kappa(\boldsymbol{\mu}\cdot\boldsymbol{n})^2}, \tag{A20}$$

where M is the first type confluent hypergeometric function [10] and is also referred to as Kummer function. Here, $\boldsymbol{\mu}$, $\kappa$ and $\boldsymbol{n}$ are denoted as the mean orientation of the Watson distribution, concentration parameter, and the orientation of sticks in which water diffusion is restricted, respectively. Since the Watson distribution is also a function of $\boldsymbol{\mu}$ and $\kappa$, these variables are expressed as $f(\boldsymbol{n}) = f(\boldsymbol{n}|\boldsymbol{\mu}, \kappa)$.

Let $\boldsymbol{\mu} = \hat{\boldsymbol{z}}$ (unit vector in the z direction) and let $x = cos\theta$, $dx = -sin\theta \cdot d\theta$, we integrate over unit sphere $\mathbb{S}^2$.

$$\int_{\mathbb{S}^2} f(\boldsymbol{n}|\hat{\boldsymbol{z}}, \boldsymbol{\kappa}) d\boldsymbol{n} = \frac{1}{M\left(\frac{1}{2}, \frac{3}{2}, \kappa\right)} \int_{\mathbb{S}^2} e^{\kappa(\hat{\boldsymbol{z}}\cdot\boldsymbol{n})^2} d\boldsymbol{n}$$

$$= \frac{1}{M\left(\frac{1}{2}, \frac{3}{2}, \kappa\right)} \int_0^{2\pi} sin\theta d\phi \int_0^{\pi} d\theta \cdot e^{\kappa(cos\theta)^2}$$

$$= \frac{1}{M\left(\frac{1}{2}, \frac{3}{2}, \kappa\right)} \cdot 2\pi \cdot \int_{-1}^{1} e^{\kappa x^2} dx\,. \tag{A21}$$

According to Arfken and Wever (2005),

$$\int_{-1}^{1} P_l(\mu)\, e^{x\mu^2} = (x)^{l/2} \frac{\Gamma\left(\frac{l+1}{2}\right)}{\Gamma\left(\frac{2l+3}{2}\right)} M\left(\frac{l+1}{2}, \frac{2l+3}{2}, x\right), \tag{A22}$$

where $\Gamma(x)$ is Gamma function, $\Gamma(1/2) = \sqrt{\pi}$, $\Gamma(3/2) = \sqrt{\pi}/2$.

Hence, Eq. A21 is expressed using Eq. (A22) as follows:

$$\int_{\mathbb{S}^2} f(\boldsymbol{n}|\hat{\boldsymbol{z}},\boldsymbol{\kappa})d\boldsymbol{n} = \frac{1}{M\left(\frac{1}{2},\frac{3}{2},\kappa\right)}\cdot 2\pi \cdot (\kappa)^{0/2}\frac{\Gamma\left(\frac{1}{2}\right)}{\Gamma\left(\frac{3}{2}\right)}M\left(\frac{1}{2},\frac{3}{2},\kappa\right)$$

$$= 4\pi. \qquad (A23)$$

Since we want to normalize the Watson distribution, we re-defined it as follows:

$$f(\boldsymbol{n}|\boldsymbol{\mu},\kappa) = \frac{1}{4\pi\cdot M\left(\frac{1}{2},\frac{3}{2},\kappa\right)}e^{\kappa(\boldsymbol{\mu}\cdot\boldsymbol{n})^2}. \qquad (A24)$$

3.2. *Simulation for relationship between NODDI and DTI-derived NODDI under the condition that CSF volume fraction was homogeneously fixed.*

The previous simulation (see section 2.2 and 3.2 in main text) suggested that heterogeneity in CSF volume fraction in the cortical voxels can cause the errors at various level particularly when used low b-value dMRI dataset. To confirm the specificity of this finding, we performed a simulation analysis whether fixed 'homengenous' values of $V_{iso}$ in the cortical voxels does not degrade a relationship between the original NODDI and DTI-derived NODDI depending on the b-shell scheme. The size of the CSF compartment ($V_{iso}$) in the cortex was assumed to be homogeneous and small in a simulation analysis ($V_{iso}$=0.1). Seven different combinations of b-shell datasets (same as Table 1) were created assuming following parameters as possible values within the cerebral cortex[8]; $V_{iso}$=0.1, NDI ranging from 0.1 to 0.55 and ODI ranging from 0.040 to 0.84 (see Table S1).

To investigate linearity, $NDI_{DTI}$ and $ODI_{DTI}$ were correlated with the true values using the Pearson correlation analysis for each dataset. As a result, $NDI_{DTI}$ and $ODI_{DTI}$ showed extremely strong linear correlation with the ground truth in any b-shell scheme ($R>0.97$, $p<0.00001$) (Fig. S2).

Like in HCP data (see Fig. 5 in main text), a constant bias between DTI-derived NODDI and original NODDI parameters was found in the Bland-Altman analysis, when using simulation data with high b-value. This bias was caused by the bias of the MD and FA when using high b-value dataset — the values of MD were underestimated and those of FA were overestimated (data not shown), consistent with previous studies in MD[11] and FA[12–14].

**Table S1**. Parameters and values used in simulation analysis. Note that all combinations of values of NDI and ODI were simulated.

| | | | | | | | | | | |
|---|---|---|---|---|---|---|---|---|---|---|
| NDI | 0.1 | 0.15 | 0.2 | 0.25 | 0.3 | 0.35 | 0.4 | 0.45 | 0.5 | 0.55 |
| ODI | 0.040 | 0.11 | 0.16 | 0.30 | 0.37 | 0.47 | 0.55 | 0.61 | 0.70 | 0.84 |

**Figure S1**

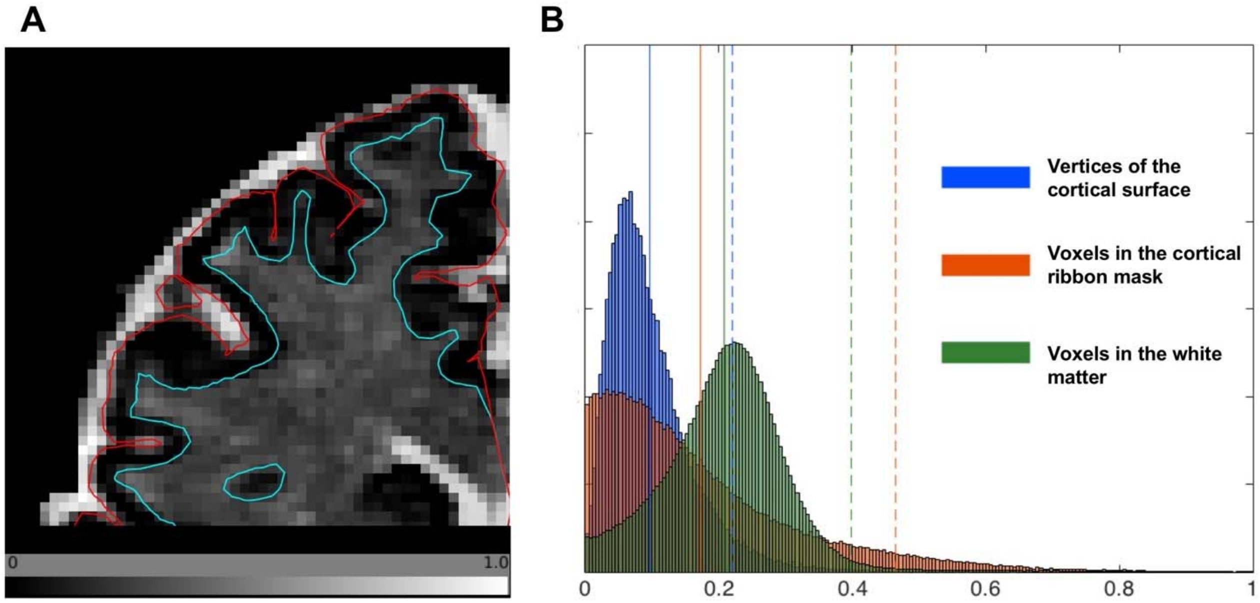


**Figure S1.** The $V_{iso}$ (volume of cerebrospinal fluid compartment) in the cerebral cortex. **(A)** The zoomed view of a representative volume slice of $V_{iso}$ in a single subject. The red line is the pial surface, which is the border between CSF and the cortex, and the blue line is the white surface, which is the border between the cortex and the white matter. The value of $V_{iso}$ in the cortex is very low compared to the white matter and CSF. **(B)** The histogram of $V_{iso}$ in the bilateral cerebral cortices in volume-based and surface-based analysis and the white matter. in the corresponding subject in **(A)**. The blue histogram shows vertices of the cortical surface. The orange histogram shows voxels in the cortical ribbon mask. The green histogram shows voxels in the white matter. Colored solid lines indicate the mean and colored dotted lines indicate the mean+1.96*SD.

**Figure S2**

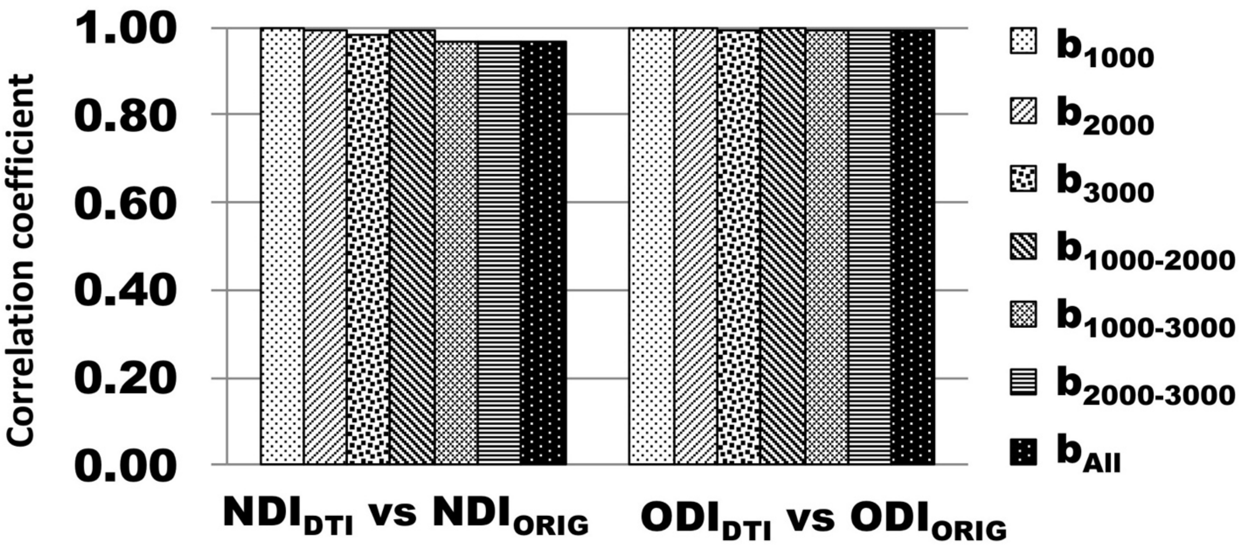


**Figure S2.** correlation coefficients of DTI-derived NODDI parameters ($NDI_{DTI}$ and $ODI_{DTI}$) with respect to the ground truth in simulation analysis. Correlation coefficients were calculated using various b-shell dataset types ($b_{1000}$, $b_{2000}$, $b_{3000}$, $b_{1000-2000}$, $b_{1000-3000}$, $b_{2000-3000}$ and $b_{All}$) . All of them have statistical significance level with $p<0.00001$. Note that this simulation does not consider 'heterogeneity' of CSF volume fraction (see also Figure 6 for simulation of heterogeneity of CSF). Abbreviations; $NDI_{ORIG}$: original NODDI neurite density index, $ODI_{ORIG}$: original NODDI orientation dispersion index, $NDI_{DTI}$: DTI-derived NODDI neurite density index, $ODI_{DTI}$: DTI-derived NODDI orientation dispersion index.

**Figure S3**

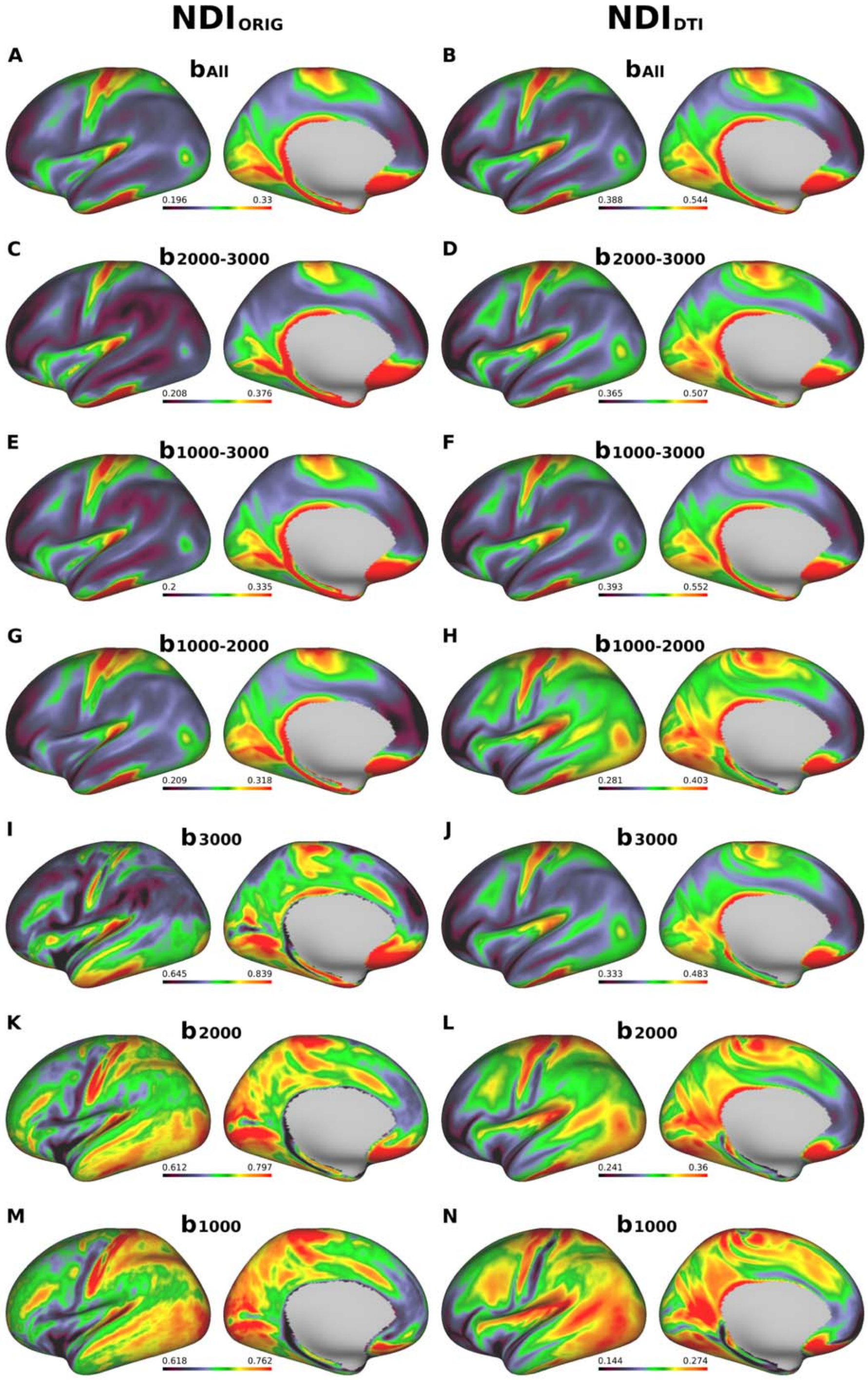


**Figure S3** Cross-subject average cortical surface maps of original NODDI and DTI-derived NODDI neurite density index ($NDI_{ORIG}$ and $NDI_{DTI}$) when using b-shell datasets ($b_{1000}$, $b_{2000}$, $b_{3000}$, $b_{1000\text{-}2000}$, $b_{1000\text{-}3000}$, $b_{2000\text{-}3000}$, $b_{All}$).
Surface maps of column 1 **(A, C, E, G, I, K, M)** were cortical maps of the original NODDI NDI ($NDI_{ORIG}$) and surface maps of column 2 **(B, D, F, H, J, L, N)** were cortical maps of DTI-derived NDI ($NDI_{DTI}$). The first through the seventh rows are arranged in order of the b-shell datasets. Data at https://balsa.wustl.edu/977qk

**Figure S4**

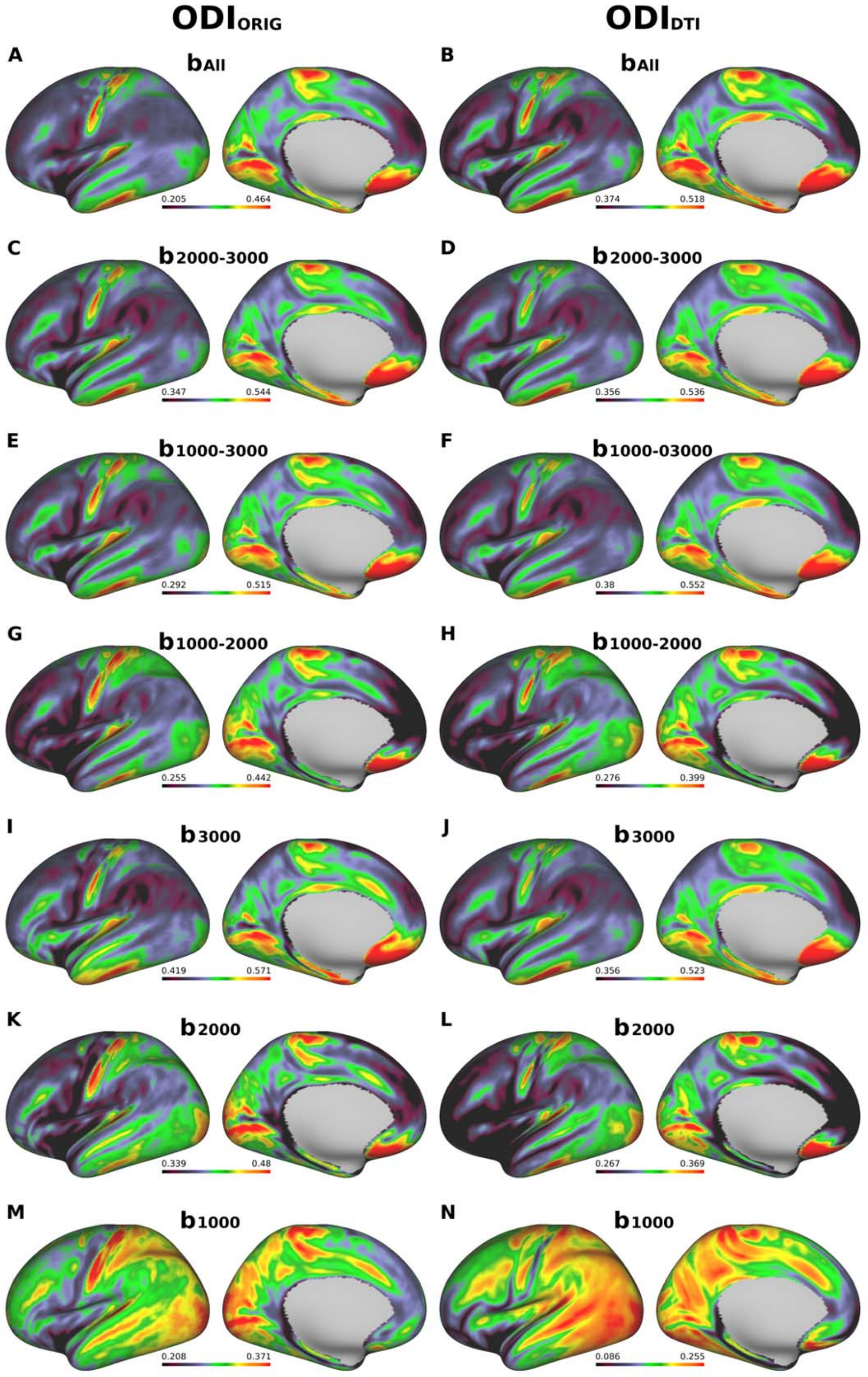


**Figure S4.** Cross-subject average cortical surface maps of original NODDI and DTI-derived NODDI orientation dispersion index ($ODI_{ORIG}$ and $ODI_{DTI}$) when using b-shell datasets ($b_{1000}$, $b_{2000}$, $b_{3000}$, $b_{1000\text{-}2000}$, $b_{1000\text{-}3000}$, $b_{2000\text{-}3000}$, $b_{All}$). Surface maps of column 1 **(A, C, E, G, I, K, M)** were cortical maps of the original NODDI ODI ($ODI_{ORIG}$) and surface maps of column 2 **(B, D, F, H, J, L, N)** were cortical maps of DTI-derived ODI ($ODI_{DTI}$). The first through the seventh rows are arranged in order of the b-shell datasets. https://balsa.wustl.edu/kNNK2

**Figure S5**

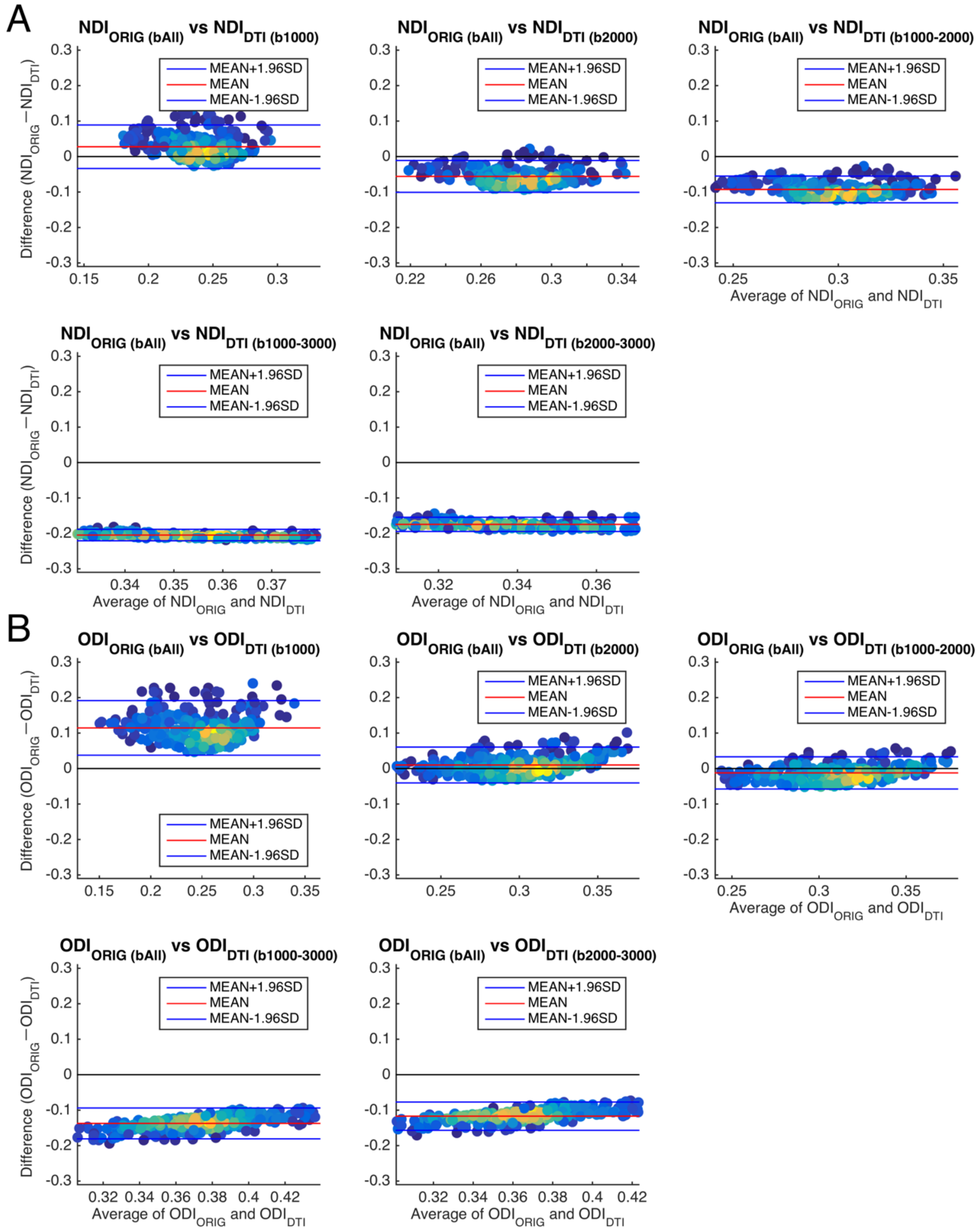


**Figure S5** Bland-Altman plots of DTI-derived NODDI parameters of different b-shell datasets different b-shell datasets ($b_{1000}$, $b_{2000}$, $b_{1000\text{-}2000}$, $b_{1000\text{-}3000}$, and $b_{2000\text{-}3000}$). Biases of DTI-derived NODDI parameters ($NDI_{DTI}$ shown in **(A)** and $ODI_{DTI}$ shown in **(B)** in different b-shell datasets ($b_{1000}$, $b_{2000}$, $b_{1000\text{-}2000}$, $b_{1000\text{-}3000}$, and $b_{2000\text{-}3000}$) were analyzed with Bland-Altman and the original NODDI parameters ($NDI_{ORIG}$ and $ODI_{ORIG}$) of the three-shell dataset ($b_{All}$) used as a reference. Blue lines show the mean±1.96*SD and red lines show the mean value. Plots are coloured by their density. Abbreviations; $NDI_{ORIG}$: original NODDI neurite density index, $ODI_{ORIG}$: original NODDI orientation dispersion index, $NDI_{DTI}$: DTI-derived NODDI neurite density index, $ODI_{DTI}$: DTI-derived NODDI orientation dispersion index.

**Figure S6**

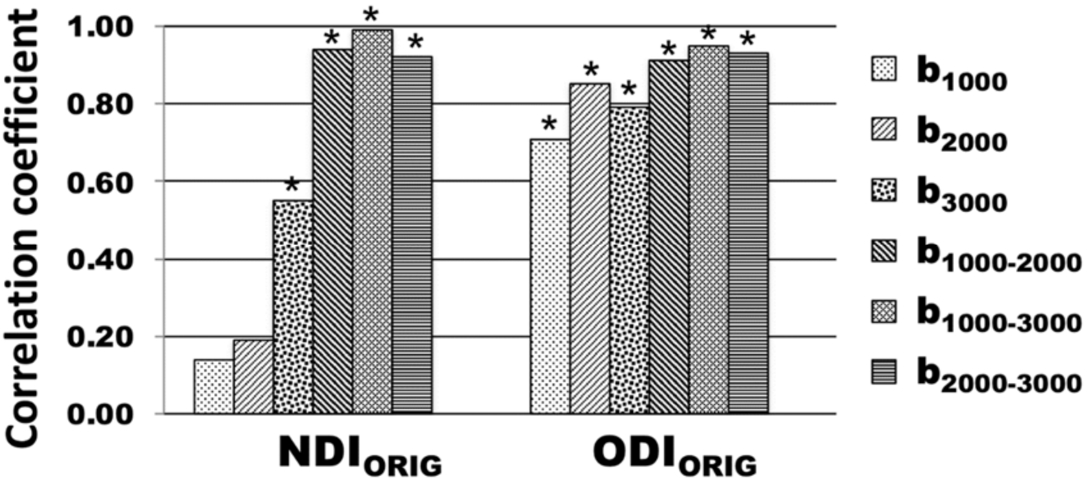


**Figure S6**. Correlation coefficients of original NODDI parameters ($NDI_{ORIG}$ and $ODI_{ORIG}$) with the references that are the original NODDI parameters on three-shell dataset ($NDI_{ORIG}/b_{All}$ and $ODI_{ORIG}/b_{All}$). Correlation coefficients to the references were calculated using averaged surface maps across all subjects in each different b-shell dataset type ($b_{1000}$, $b_{2000}$, $b_{3000}$, $b_{1000-2000}$, $b_{1000-3000}$, and $b_{2000-3000}$). Asterisks were significant ($p<0.00001$).
